# THE FAM53C/DYRK1A axis regulates the G1/S transition of the cell cycle

**DOI:** 10.1101/2024.12.10.627280

**Authors:** Taylar Hammond, Jong Bin Choi, Miles W. Membreño, Janos Demeter, Roy Ng, Debadrita Bhattacharya, Thuyen N. Nguyen, Griffin G. Hartmann, Caterina I Colón, Carine Bossard, Jan M. Skotheim, Peter K. Jackson, Anca Pasca, Seth M. Rubin, Julien Sage

## Abstract

A growing number of therapies are being developed to target the cell cycle machinery for the treatment of cancer and other human diseases. Consequently, a greater understanding of the factors regulating cell cycle progression becomes essential to help enhance the response to these new therapies. Here, using data from the Cancer Dependency Map, we identified FAM53C as a new regulator of cell cycle progression. We found that FAM53C is critical for this cell cycle transition and that it acts upstream of the CyclinD-CDK4/6-RB axis and of p53 in the regulation of the G1/S transition. By mass spectrometry, biochemical, and cellular assays, we identified and validated DYRK1A as a cell cycle kinase that is inhibited by and directly interacts with FAM53C. Consistent with the role for FAM53C identified in cells in culture, *FAM53C* knockout human cortical organoids display increased cell cycle arrest and growth defects. *Fam53C* knockout mice show minor behavioral phenotypes. Because DYRK1A dysregulation contributes to developmental disorders such as Down syndrome as well as tumorigenesis, future strategies aiming at regulating FAM53C activity may benefit a broad range of patients.

## INTRODUCTION

The cell cycle is precisely regulated by a complex network of checkpoints. Among those, the G1 checkpoint is an essential decision point for cell cycle progression into S-phase or cell cycle exit in G0, either for a transient arrest, such as quiescence, or for a more permanent arrest, such as during cellular differentiation or senescence. Signal integration at this cell cycle checkpoint is tightly regulated, and defects at this transition can have catastrophic impacts in development and human diseases, including auto-immune diseases and cancer [1]. A better understanding of the mechanisms regulating the G1/S decision point is critical for our understanding of embryonic development, tissue repair, regeneration, and aging processes.

The G1/S transition is regulated by multiple intracellular and extracellular signals, many of which converge onto the RB pathway [2]. The core RB pathway is centered around the retinoblastoma (RB) tumor suppressor. RB normally inhibits cell cycle progression in G1 by interacting with various proteins, including the E2F family of transcription factors, which regulate the expression of genes essential for DNA replication and other aspects of cell division [3]. Under the prevailing model, mitogen signaling stimulates RB hyperphosphorylation and inactivation by Cyclin/CDKs (namely Cyclin D-CDK4/6 and Cyclin E-CDK2), leading to E2F release to promote proliferation; mechanisms such as upregulation of the p21 cell cycle inhibitor (p21WAF1/CIP1, encoded by the *CDKN1A* gene) under stress conditions, often downstream of p53 activation [4], can limit CDK activity to keep cells in G1 arrest [4].

Cyclin D (with three family members Cyclin D1, D2, and D3), is a critical Cyclin whose levels are tightly regulated both at the transcriptional level and the post-transcriptional level, including by regulation of protein stability to induce rapid degradation [5, 6]. Recently, we and others have shown that AMBRA1 (Activating Molecule in Beclin1-Regulated Autophagy) is an adaptor for the CRL4 (CUL4-RING) E3 ubiquitin ligase to mediate the polyubiquitylation and subsequent degradation of cyclin D proteins in cells [7–9]. These studies resolved a long-standing question in the field related to the molecular mechanisms regulating Cyclin protein levels regulation, but also highlighted that, despite decades of research, key regulators of core cell cycle machinery remain to be discovered and characterized.

Accumulating evidence also indicates that Dual-specificity Tyrosine Phosphorylation-regulated Kinase 1A (DYRK1A), and its family member DYRK1B, can phosphorylate Cyclin D (at threonine 286, T286, for Cyclin D1), which leads to lower Cyclin D protein levels in cells [10–16] while also modulating p21 levels in a bistable manner [10]. DYRK1A has gained clinical interest due to its dosage sensitive role in cancer and developmental disease: the *DYRK1A* gene maps to the Down syndrome critical region on chromosome 21q22 [17, 18] and increased DYRK1A kinase activity has been linked to many of the developmental brain defects associated with Down syndrome [19–21]. In addition, functional loss of one *DYRK1A* allele leads to the so-called DYRK1A haploinsufficiency syndrome [22]. *DYRK1A* variants have also recently been associated with epilepsy [23]. The regulation of DYRK1A and its effects toward Cyclin D1 and p21 in normal and disease contexts remain poorly understood.

Here we aimed to identify new regulators of the G1/S transition of the cell cycle. Using co-dependency data in hundreds of human cancer cell lines from the Cancer Dependency Map Project (DepMap, [24]), we identified the poorly studied protein FAM53C as a candidate cell cycle regulator connected to the RB pathway. Our work shows that FAM53C normally promotes G1/S progression and links its cell cycle activity to DYRK1A activity in cells.

## MAIN TEXT

### A DepMap analysis identifies FAM53C as a candidate regulator of G1/S

To identify new candidate regulators of the G1/S switch, we took advantage of the co-dependency scores offered by the Cancer Dependency Map (DepMap) platform to identify potential novel pathway interactors [25, 26]. To find novel interactions, we first generated a list of 38 factors playing a known role at this cell cycle transition (**Table S1**).

We next overlapped the top 100 co-dependencies from the Cancer Dependency Map (DepMap) platform [27] for each of the input genes to generate a list of shared co-dependency genes from across the input list (**Figure 1A** and **Table S2**). We established a score for each gene hit based on the number of input genes which shared a co-dependency with the hit. The gene coding for CDK2 had the maximum of 13 hits in the analysis, while genes coding for other established G1/S factors such as E2F1 or Cyclin E2 had a score of 3, establishing a cutoff for new candidates; this cutoff score encompassed approximately the top 5% of output values (**Figure 1B,C** and **Table S2**).

**Figure 1:**
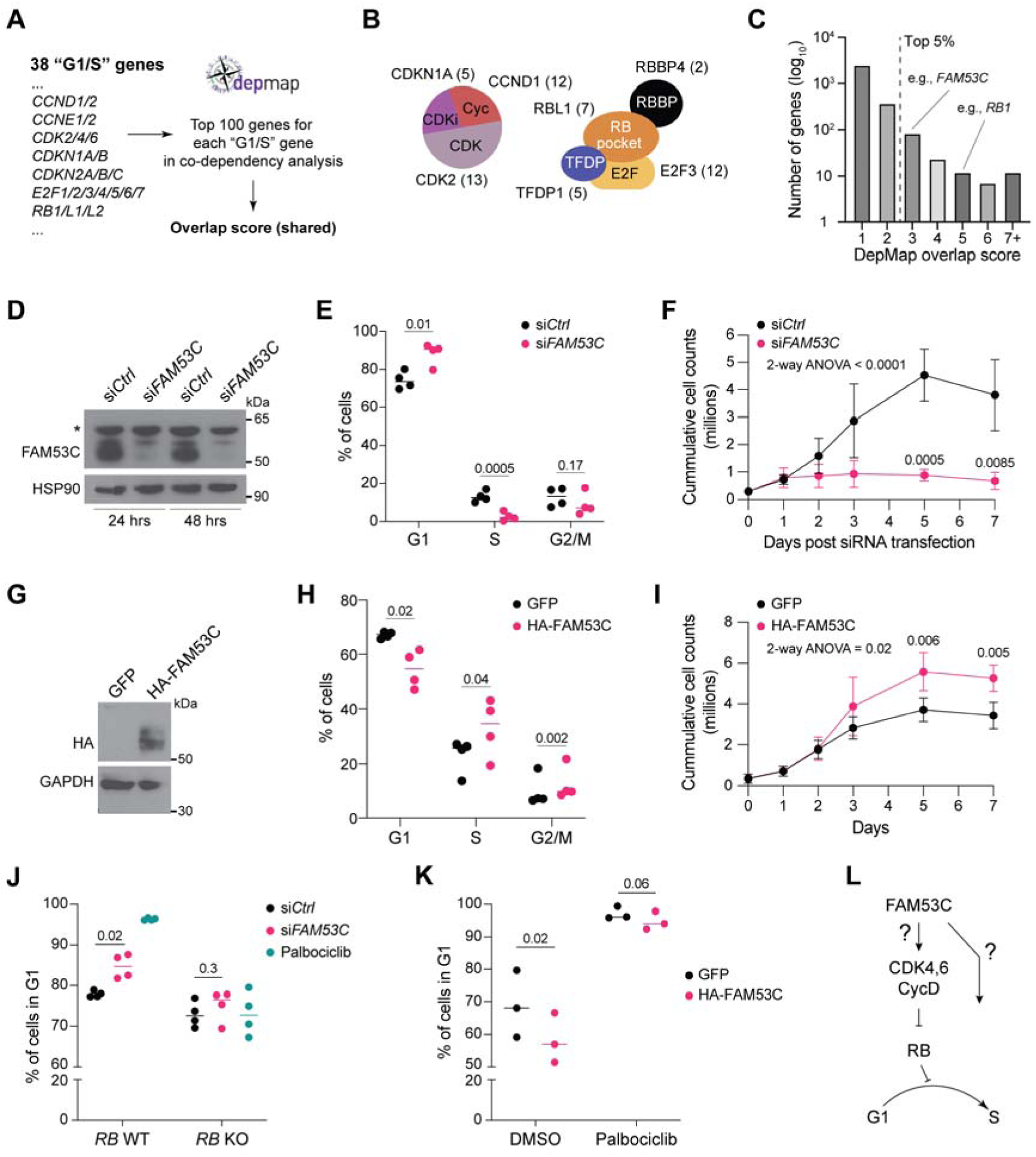
Identification of FAM53C as a positive regulator of cell cycle progression in G1. **A.** Strategy to identify new cell cycle regulators using the DepMap database. See Supplementary Table S1 for the complete list of the 38 genes and associated DepMap data. **B.** Schematic representation of key factors in the G1/S machinery and scores in the screen for selected factors. CDKi, CDK inhibitors; RBBP, RB binding protein. **C.** Representation of the number of genes with different overlap scores in the DepMap analysis. *RB1* and *FAM53C* are indicated; the maximal overlap score was 13 for the *CDK2* (not shown on graph). The cut-off overlap score value for top candidates was set at 3 (top 5%). See Supplementary Table S2 for complete data. **D.** Immunoblot analysis for FAM53C in knock-down (si*FAM53C*) RPE-1 cells compared to controls (si*Ctrl*) at 24 h and 48 h. HSP90 serves as a loading control. Molecular weights are indicated in kDa. *, non-specific signal. **E.** Cell cycle analysis by BrdU/PI staining in control and FAM53C knock-down RPE-1 cells (N=4). **F.** Population growth analysis (cell counts) in control and FAM53C knock-down RPE-1 cells (N=3-5). **G.** Representative (n=3) immunoblot analysis for FAM53C overexpression (HA-tagged FAM53C) compared to control cells (with GFP expression) in RPE-1 cells. GAPDH serves as a loading control. Molecular weights are indicated in kDa. **H.** Cell cycle analysis by BrdU/PI staining in control and FAM53C-overexpressing RPE-1 cells (n=4). **I.** Population growth analysis in control and FAM53C-overexpressing RPE-=1 cells (n=3-5). **J.** Fraction of cells in G1 from BrdU/PI staining in control (wild-type, WT) and RB knockout (KO) RPE-1 cells, with or without FAM53C knock-down (N=4). **K.** Fraction of cells in G1 from BrdU/PI staining in control (GFP) and FAM53C-overexpressing RPE-1 cells, with or without palbociclib treatment (N=3). **L.** Cartoon placing FAM53C in the G1/S transition of the cell cycle. P-values for (EH), (J), and (K) were calculated by paired t-test. P-values for (F) and (I) were calculated by mixed-model ANOVA followed by post-hoc paired two-tail t-test.

Among the genes with a score greater or equal to 3 and with no obvious prior direct link to G1 and the RB pathway, we noted *ZZZ3* (score of 6, whose product is a subunit of the Ada-two-A-containing, or ATAC histone acetyltransferase complex [28]) and C1ORF109 (score of 5, whose product is involved in replication stress [29]) (**Table S2**). We did not pursue these genes but focused on *FAM53C* (score of 3, Family with sequence similarity 53 C, previously also known as *C5ORF6*) because a similar analysis of DepMap data using a different computational approach also identified this gene as a candidate G1/S regulator [30], and because little is known about FAM53C function.

### FAM53C promotes the G1/S transition of the cell cycle

*FAM53C* has a positive co-dependency correlation with *CCND1* and *CDK4*, and negative correlations with *RB1* in the top 100 co-dependencies for each of the 38 selected factors (**Table S2**), suggesting that FAM53C may normally act as a promoter of cell cycle progression in G1. Indeed, when we acutely knocked-down FAM53C in immortalized human RPE-1 cells using short interfering RNAs (siRNAs) (**Figure 1D**), we observed a significant accumulation of cells in G1/S and loss of S-phase representation in BrdU/PI assays (**Figure 1E** and **Figure S1A** and **S1B**). This growth defect was sustained, with no population doublings over 7 days post-transfection (**Figure 1F**). No apoptosis was identified by Annexin V/PI FACS staining, confirming that loss of replication is due to true cell cycle arrest and not cell death (**Supplementary Figure S1C,D**). Knock-down of FAM53C in U2OS osteosarcoma and A549 lung cancer cell lines also led to significant G1 arrest (**Figure S1E**). Conversely, upon FAM53C overexpression (**Figure 1G**), we observed a greater fraction of cells in S-phase and increased number of cells at confluency (**Figure 1H,I** and **Figure S1F**).

Based on the DepMap co-dependency analysis and the knock-down data, we next decided to investigate the pathway interplay between the Cyclin D1/CDK4-RB pathway and FAM53C. We found that knockout of RB in RPE-1 cells significantly abrogated the G1 arrest observed upon FAM53C knock-down; loss of RB also abrogated the effects of the CDK4/6 inhibitor palbociclib in these experiments, as expected (**Figure 1J** and **Supplementary Figure S1H**). The increased proliferation of RPE-1 cells overexpressing FAM53C could be effectively blocked by palbociclib treatment (**Figure 1K** and **Supplementary Figure S1I**). Altogether, these analyses and functional data identified FAM53C as a regulator of the G1/S transition in immortalized and cancer cells (**Figure 1L**).

### The FAM53C interactome highlights the cell cycle regulatory role of FAM53C

FAM53C remains a largely uncharacterized protein with no known functional domains [31]. To gain further insights into the mechanisms of action of FAM53C in cells and how FAM53C may connect with the RB pathway, we analyzed the FAM53C interactome in cells. To this end, we expressed a GFP- and S-tagged (localization and affinity purification tag, LAP-tag) lentiviral FAM53C construct in 293T cells, pulled down the FAM53C protein, and performed mass spectrometry on the purified fraction (**Figure 2A** and **Figure S2A**). This dual-affinity purification/mass spectroscopy (AP-MS) analysis revealed candidate interactors (**Table S3**), with an enrichment for cell cycle factors (**Table S4**). Candidate FAM53C interactors in the cell cycle were related to both the G1/S (e.g., CDK4) and the G2/M (e.g., PLK1) transitions, including binding to several subunit of protein phosphatase 2A (PP2A) and several members of the 14-3-3 family, which may contribute to regulation at multiple cell cycle transitions [32, 33] (**Figure 2B** and **Table S3**). This analysis further points to a role for FAM53C at the G1/S transition of the cell cycle, but also suggests that FAM53C could play some yet unidentified roles at other phases of the cell cycle. While a specific analysis of the FAM53C interactome has not been performed previously, FAM53C was identified in other pull-down experiments. Integrating our AP/MS data with interactome data from public databases confirmed a likely role for FAM53C in the regulation of cell cycle progression, including the G1/S transition (**Figure 2C**).

**Figure 2:**
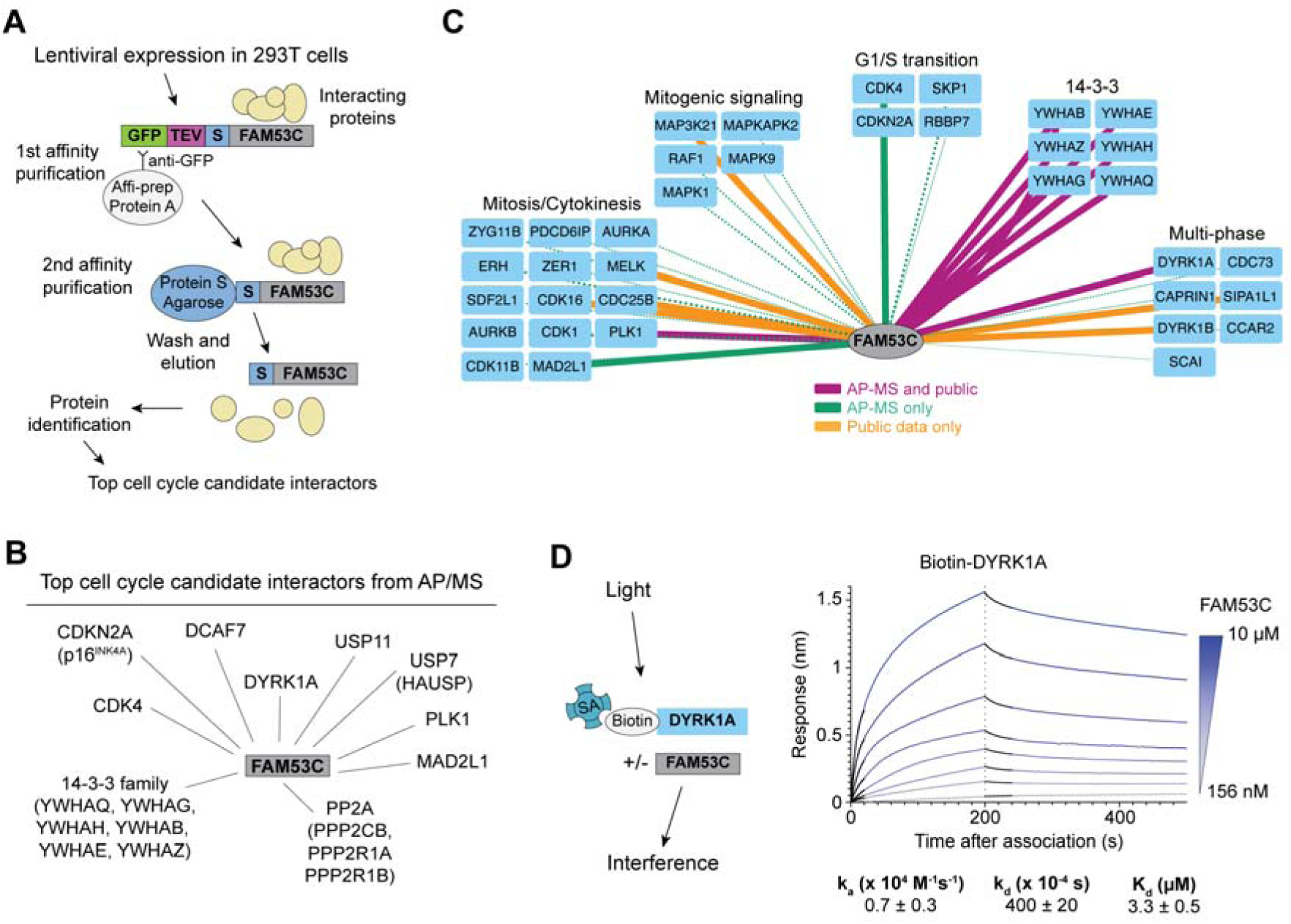
The FAM53C interactome identifies cell cycle factors. **A.** Cartoon representation of the AP/MS experiment to determine the FAM53C interactome. **B.** Top cell cycle interactors from the AP/MS experiment. See Supplementary Tables S3 and S4 for the complete list. **C.** CytoTRACE analysis integrating the results of the AP-MS experiment with public databases. Solid lines indicate p ≤ 0.05, and line thickness correlates to pNSAF from AP-MS experiment. **D.** Biolayer interferometry assay to measure the binding of recombinant FAM53C to DYRK1A-coated streptavidin (SA) sensors. Association begins a 0 seconds. Dotted line indicates start of dissociation phase. Black solid line indicates portion of the curve that was analyzed for on and off rates. Reported rate constants are from data fitting of observed on and off rates and the equilibrium constant was determined from the steady-state response analysis.

While FAM53C is likely to have several important partners in cells, DYRK1A stood out as a promising candidate. First, previous proteomic studies focusing on the DYRK1A interactome had identified FAM53C in the list of potential partners of DYRK1A [34–37]. Second, our proteomic analysis also identified DCAF7, which is a known functional partner of DYRK1A [38, 39]. Third, a recent study indicates a functional relationship between FAM53C and DYRK1A in neurons [40]. Fourth, strong evidence indicates that DYRK1A kinase activity is a key regulator of the G1/S transition: DYRK1A phosphorylation of LIN52 can promote the activity of the Dimerization Partner, RB-like, E2F, and multivulva class B (DREAM) complex, thereby promoting quiescence [41]; DYRK1A phosphorylation of Cyclin D1 at T286 also results in Cyclin D1 degradation, thereby slowing or preventing G1/S progression [10, 12–14, 42, 43]. Thus, to further assess a potential binding interaction between FAM53C and DYRK1A, we used biolayer interferometry (BLI), an optical biosensing technology that quantifies molecular interactions. After affinity purification of FAM53C and DYRK1A expressed from bacteria, a sortase reaction was used to label the N-terminus of DYRK1A with a biotin tag, which allowed us to efficiently load DYRK1A onto streptavidin-coated BLI sensors. FAM53C association to the sensors was then measured in a dose-dependent manner, and rate and binding constants were determined (**Figure 2D**). Using this approach, we found that FAM53C has an affinity for DYRK1A of K_d_ = 3.3 ± 0.5 μM, confirming the ability of these two proteins to directly interact. Based on these observations, we decided to pursue the analysis of the interactions between FAM53C and DYRK1A in cells.

### FAM53C acts as an inhibitor of DYRK1A

The direct binding of FAM53C to DYRK1A and the co-dependency scores between FAM53C, DYRK1A, and Cyclin D1 in the DepMap analysis led us to hypothesize that FAM53C may act as a direct inhibitor of DYRK1A kinase activity, in addition to its recently reported role as a regulator of DYRK1A localization in neurons [40]. To test this idea, we first developed a radio-labeling kinase assay utilizing recombinantly expressed DYRK1A, Cyclin D1, and FAM53C. Cyclin D1 phosphorylation by DYRK1A has been previously observed *in vitro* at short time points (<30 min) [10, 12–14, 42, 43], making it a biochemically and biologically relevant substrate to test FAM53C effects on DYRK1A. Upon titrating increasing amounts of FAM53C, we observed a reduction in Cyclin D1 phosphorylation in this assay (**Figure 3A**). We also noted efficient phosphorylation of FAM53C itself in the presence of DYRK1A with lower levels of phosphorylation at lower levels of FAM53C (**Figure 3A**), suggesting that FAM53C may be a competitive substrate and/or an inhibitor of DYRK1A. Phosphorylation of LIN52, another substrate of DYRK1A [41] was also reduced with increasing levels of FAM53C (**Figure S3A**). Taken together these data support a model in which FAM53C can inhibit DYRK1A kinase activity toward key cell-cycle substrates.

**Figure 3:**
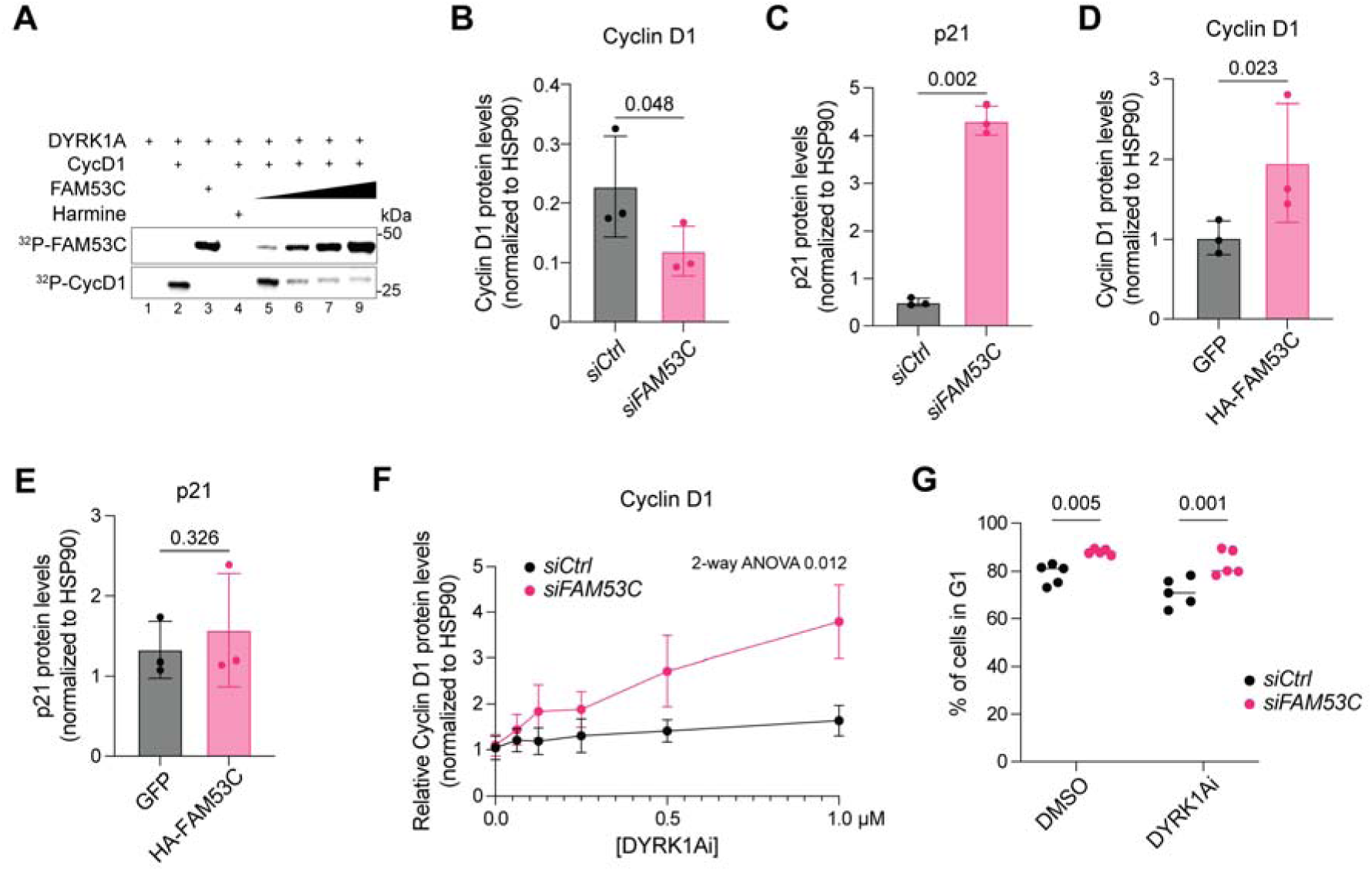
FAM53C can inhibit DYRK1A function in cells. **A.** *In vitro* phosphorylation assay. Recombinantly expressed Cyclin D1 (1 µM) was phosphorylated by DYRK1A (200 nM) for 15 min alone or in the presence of increasing amounts of FAM53C. FAM53C concentration was 5 µM (lane #3) or 1, 2.5, 5, and 10 µM (lanes #5-8). Lane #1 contains DYRK1A without substrate. The DYRK1A kinase inhibitor Harmine was added at 25 µM. Representative experiment of N=3 experiments. **B.** Quantification of immunoassays for Cyclin D1 protein levels relative to the loading control HSP90 in FAM53C knock-down (si*FAM53C*) RPE-1 cells compared to controls (si*Ctrl*) at 24 h (N=3). **C.** Quantification of immunoassays for p21 protein levels relative to the loading control HSP90 in FAM53C knock-down (si*FAM53C*) RPE-1 cells compared to controls (si*Ctrl*) at 24 h (N=3). **D.** Quantification of immunoassays for Cyclin D1 protein levels relative to the loading control HSP90 in RPE-1 cells expressing HA-FAM53C compared to GFP controls (N=3). **E.** Quantification of immunoassays for p21 protein levels relative to the loading control HSP90 in RPE-1 cells expressing HA-FAM53C compared to GFP controls (N=3). **F.** Quantification of immunoassays for Cyclin D1 protein levels in control and FAM53C knock-down RPE-1 cells treated with or without different concentrations of the SM13797 DYRK1Ai (N=3 per concentration, 48 hours of knock-down and treatment). **G.** Fraction of FAM53C knock-down (si*FAM53C*) RPE-1 cells in G1 compared to controls (si*Ctrl*), with or without DYRK1Ai treatment (N=5). P-values for (B), (C), (D), and (E) were calculated by paired t-test. P-value for (F) was calculated by 2-way ANOVA.

DYRK1A dosage has been linked to shifting dynamics between the cell cycle regulators Cyclin D1 and p21, including a cell cycle arrest state characterized by low levels of Cyclin D1 and high levels of p21 associated with increased DYRK1A activity [10]. When we examined Cyclin D1 and p21 protein levels upon FAM53C knock-down, we found that Cyclin D1 levels decreased while p21 levels increased (**Figure 3B,C** and **Figure S3B,C**), consistent with increased DYRK1A activity upon FAM53C loss. Overexpression of FAM53C led to increased levels of Cyclin D1, with no changes on p21 levels (**Figure 3D,E** and **Figure S3D,E**). These results further support a model in which FAM53C acts as a DYRK1A inhibitor. These data also raised the possibility that FAM53C levels and p21 levels may not be only linked by DYRK1A activity.

To test whether DYRK1A inhibition could rescue the G1 arrest observed upon *FAM53C* knock-down, we used a highly selective ATP-competitive DYRK1A inhibitor (SM13797 [44], hereafter DYRK1Ai) (**Figure S3F-H**). Notably, addition of DYRK1Ai to *FAM53C* knock-down RPE-1 cells was more effective at increasing Cyclin D1 levels than in control cells (**Figure 3F** and **Figure S3I**), further supporting the model in which FAM53C acts as an active site competitor. However, DYRK1Ai treatment was insufficient to rescue the G1 accumulation induced by FAM53C loss in this context (**Figure 3G**).

Altogether, these experiments place FAM53C as an inhibitor of DYRK1A and a regulator of Cyclin D1 levels. Nevertheless, the lack of rescue of the cell cycle arrest by DYRK1Ai upon FAM53C knock-down suggested that additional mechanisms may be at play in the regulation of cell cycle progression. Notably, we found that p21 levels, which are elevated upon *FAM53C* knock-down in RPE-1 cells, were further elevated in cells treated with increasing doses of DYRK1Ai (**Figure S3J,K**). This observation led us to investigate cell cycle arrest mechanisms possibly related to p21 levels downstream of FAM53C knock-down.

### Activation of p53 downstream of FAM53C loss

The gene coding for p21, *CDKN1A*, is known to be regulated at the transcriptional level, including by p53. To determine whether there is transcriptional activation of *CDKN1A* in *FAM53C* knockdown cells, we analyzed the transcriptome of RPE-1 cells 48 hours post-knockdown. Compared to controls, FAM53C knockdown cells exhibited enriched downregulation of genes related to cell cycle processes (**Figure S4A** and **Table S5**), along with significant changes in genes coding for key cell cycle factors (**Figure S4B**), as would be expected for cells undergoing G1 arrest. Notably, a number of p53 transcriptional targets were significantly upregulated in the knock-down condition, with *CDKN1A* showing the strongest upregulation in the FAM53C knockdown cells (**Figure 4A**) and showing decreased levels in p53 knockout cells (**Figure 4B**). However, p21 knock-down, even with DYRK1Ai treatment, did not rescue the cell cycle arrest observed in FAM53C knock-down RPE-1 cells, indicating that other p53 targets are implicated (**Figure 4C** and **Figure S4C,D**).

**Figure 4:**
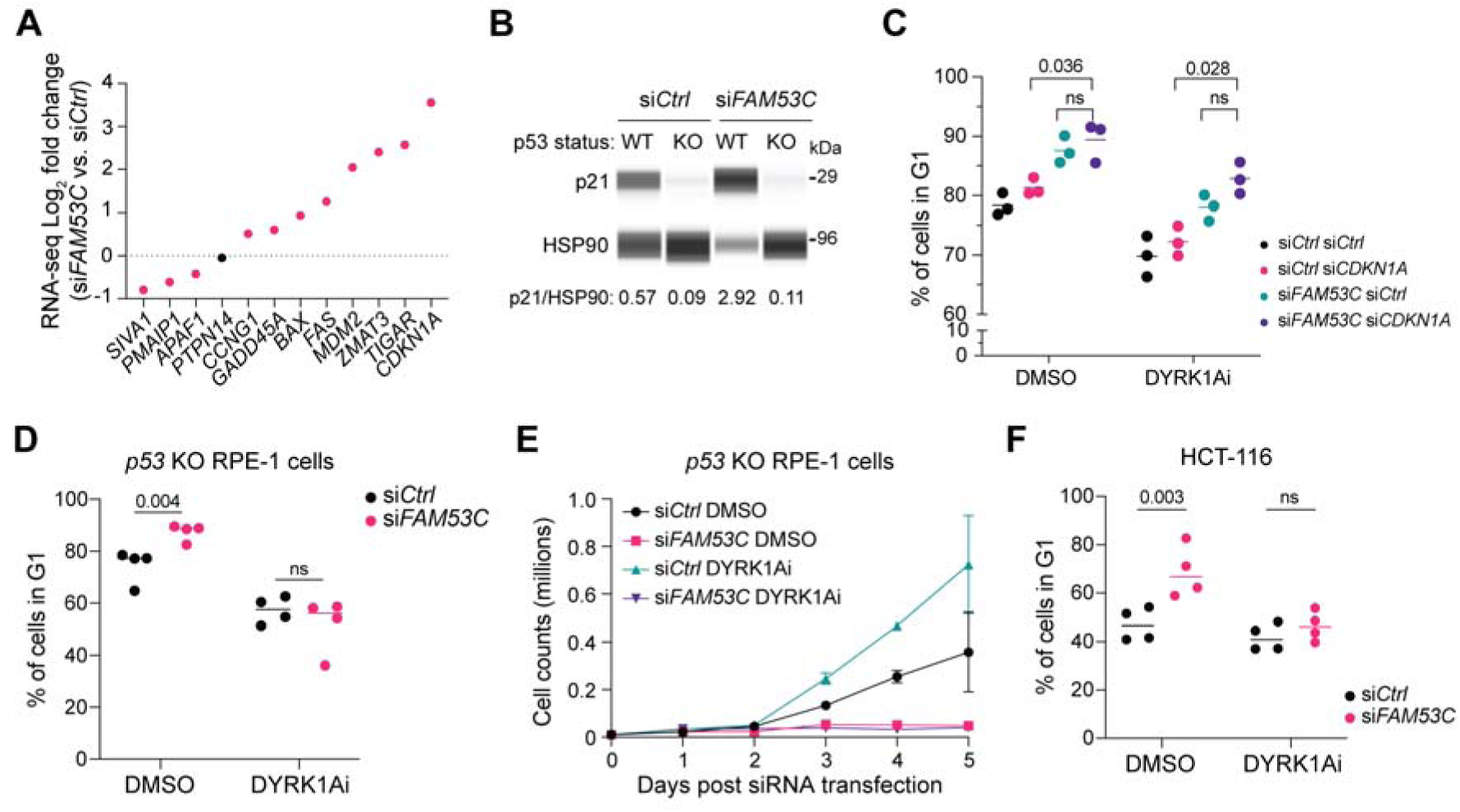
FAM53C knock-down activates p53. **A.** Fold-change analysis of the p53 target genes in FAM53C knock-down cells compared to controls (RNA-seq data). Black dot: P-value>0.05. **B.** Immunoassay for p21 in *p53* wild-type (WT) and knockout (KO) RPE-1 cells, with (si*FAM53C*) or without (si*Ctrl*) FAM53C knock-down. HSP90 serves as a loading control (N=1). Values on the bottom represent the ratio between the signal for p21 and the signal for HSP90 in the same lane. **C.** Fraction of cells in G1 in RPE-1 cells, with or without FAM53C knock-down, with or without p21 knock-down (p21 is encoded by *CDKN2A*), with or without treatment with the DYRK1A inhibitor (DYRK1Ai) (N=3). DMSO is a control for DYRK1Ai. Note that the G1 arrest observed upon FAM53C knock-down is still present in cells with p21 knock-down and DYRK1Ai treatment (ns, not significant). **D.** Fraction of FAM53C knock-down (si*FAM53C*) *p53* knockout RPE-1 cells in G1 compared to controls (si*Ctrl*), with or without DYRK1Ai treatment (N=4). Note that the G1 arrest observed upon FAM53C knock-down is absent in cells with *p53* knockout and DYRK1Ai treatment. **E.** Cell counts for *p53* knockout RPE-1 cells, with or without FAM53C knock-down, with or without DYRK1Ai treatment (N=2). Note that cells with FAM53C knock-down and DYRK1Ai treatment do not proliferate despite the decrease in G1 fraction in (D). **F.** Fraction of FAM53C knock-down (si*FAM53C*) HCT-116 cells in G1 compared to controls (si*Ctrl*), with or without DYRK1Ai treatment (N=4). Note that the G1 arrest observed upon FAM53C knock-down is absent in cells with *p53* knockout and DYRK1Ai treatment. P-values for (C), (D), and (F) were calculated by paired t-test.

Based on these observations, we next compared the effects of the of the FAM53C knockdown on the cell cycle of wild-type and *p53* knockout RPE-1 cells. Similar to DYRK1A inhibition, ablation of p53 alone was not sufficient to rescue the arrest induced by loss of FAM53C (**Figure S4E-H**, see **Figure 3G**). In contrast, the combination of *p53* knockout and treatment with the DYRK1Ai rescued G1 accumulation in FAM53C knock-down cells (**Figure 4D** and **Figure S4E**), indicating that both pathways contribute to this arrest in RPE-1 cells.

We noted that in the G1 rescue condition with DYRK1A inhibition and *p53* knockout in RPE-1 cells, S-phase values remained largely unchanged while a secondary accumulation of cells in G2/M appeared (**Figure S4E**). This observation suggested that release from *FAM53C* knockdown-mediated G1 arrest may result in stress at later phases of the cell cycle in this cell line. Indeed, when we conducted a cell count assay in *p53* knockout RPE-1 cells, cells treated with control siRNAs grew as expected, but we did not observe increased counts for FAM53C knock-down cells treated with DYRK1Ai over the course of the assay (**Figure 4E**), indicating that these cells, while entering S-phase, are not returning to a normal cell cycle. Cell lysates from cells collected at 48 hours showed an upregulation of cleaved caspase 3 (CC3) only in DYRK1Ai-treated FAM53C knock-down cells lacking p53 (**Figure S4I**), suggesting that bypass of FAM53C-loss arrest may lead to significant cell stress in later stages of the cell cycle and death.

In the DepMap dataset, p53 restricts the expansion of RPE-1-derived cell lines and Cyclin D1 is critical for their expansion. We also found that HCT-116 colon cancer cells (which are p53 wild-type) are dependent on Cyclin D1 for their expansion, similar to RPE-1 cells, but largely independent on p53 (**Figure S4J**). Notably, FAM53C knock-down in these cells led to G1 arrest, which was rescued by treatment with the DYRK1A inhibitor (**Figure 4F** and **Figure S4K**), indicating that DYRK1A is a critical mediator of cell cycle arrest in HCT-116 cells upon FAM53C knock-down and that loss of p53 is not always required to rescue the G1 arrest observed upon FAM53C knock-down. We did not examine the long-term proliferative potential of these cells, but DYRK1A inhibition has been previously shown to negatively affect the G2/M program in these cells [45]. Overall, these experiments identify activation of the p53 pathway and other stress signals downstream of FAM53C loss, in addition to activation of DYRK1A (**Figure S4L**).

### Consequences of *FAM53C* inactivation in human cortical organoids in culture

Based on the dosage-dependent role of DYRK1A in brain development, we wondered if loss of FAM53C, which can lead to DYRK1A kinase activation, may result in brain developmental phenotypes. As a first way to test this idea, we knocked out *FAM53C* in human induced pluripotent stem cells (iPSCs) using CRISPR/Cas9 (**Figure 5A** and **Figure S5A-C**). The *FAM53C* knockout in iPSCs was compatible with their survival and their stemness (**Figure S5D,E**), which allowed us to differentiate control and knockout cells into human cortical organoids (hCOs). After 25 days of differentiation, we noted that FAM53C mutant organoids were smaller in size compared to controls at this stage (**Figure 5B,C**). Using EdU incorporation assays, we also observed decreased proliferation in the knockout hCOs compared to controls (**Figure 5D** and **Figure S5F**). Loss of FAM53C in hCOs did not affect DYRK1A levels but the ratio between phosphorylated Cyclin and total Cyclin D1 levels was greater in knockout hCOs compared to controls, and p21 levels were elevated (**Figure 5F-H**). While a complete analysis of the role of FAM53C in the development of various cell types and structures in hCOs remains to be performed, these observations further support a role for FAM53C in the control of cell cycle progression in G1 in a neural development context.

**Figure 5:**
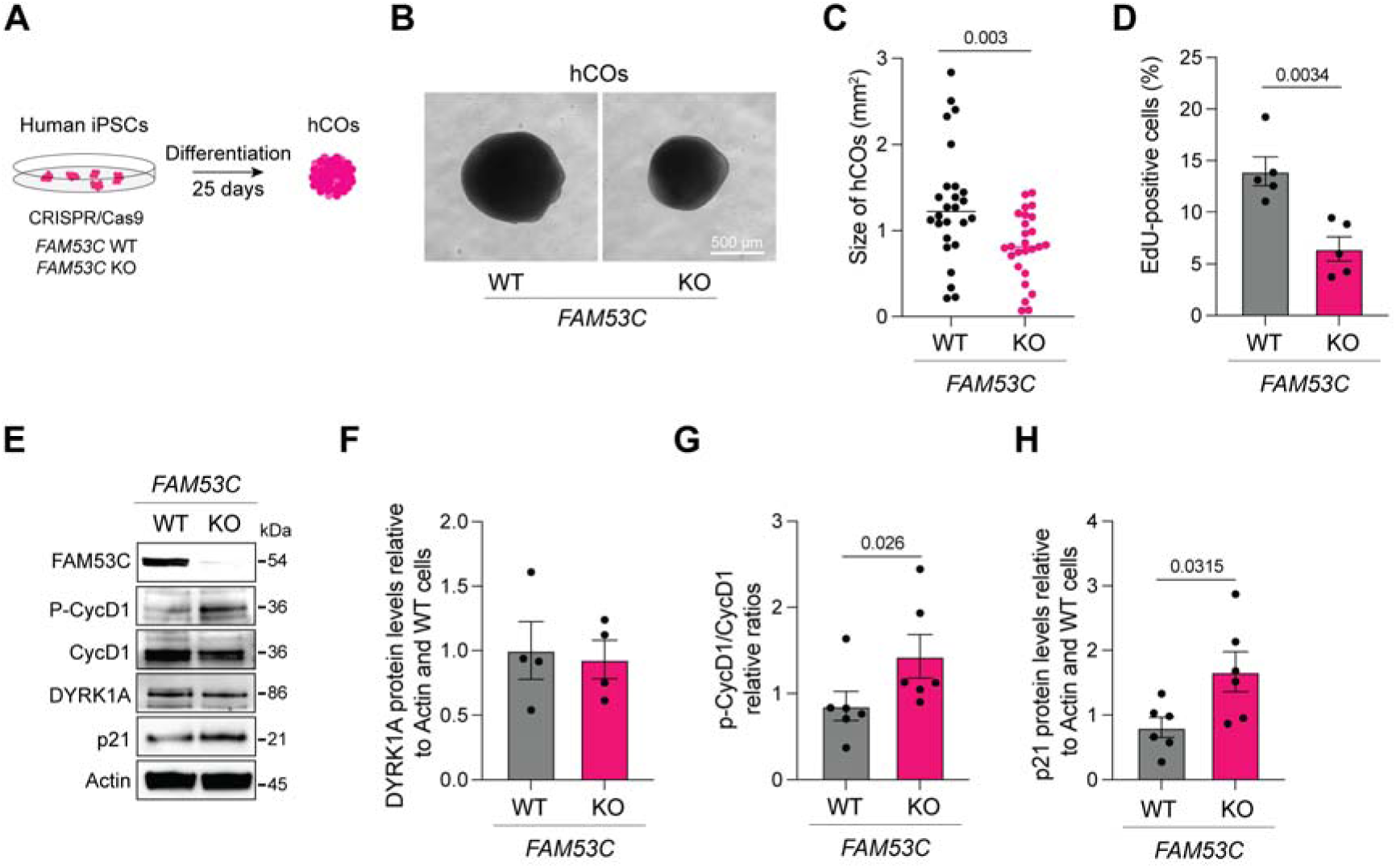
Loss of FAM53C impairs the development of human cortical organoids. **A.** Cartoon of the differentiation protocol from human induced pluripotent stem cells (iPSCs) to human cortical organoids (hCOs). **B.** Representative images of wild-type and knockout hCOs. Scale bar, 500 µm. **C.** Quantification of (B). **D.** Quantification of Edu-positive cells in wild-type and knockout hCOs. **E.** Representative immunoblot analysis of wild-type (WT) and *FAM53C* knockout (KO) hCOs. β-actin serves as a loading control. **F.** Quantification of (E) for DYRK1A relative to β-actin. **G.** Quantification of (E) for phospho-Cyclin D1 relative to Cyclin D1 levels. **H.** Quantification of (E) for p21 levels relative to β-actin levels. P-values for (C), (D), (F), (G), and (H) calculated by paired t-test.

### *Fam53C* knockout mice are viable and display only mild phenotypes

These data, and developmental roles for DYRK1A also in the brain and outside of the brain [13, 14, 46] led us to investigate the possible phenotypes of *Fam53C* knockout mice. We obtained mice with a deletion of *Fam53C* exon 4 from the International Mouse Phenotyping Consortium (IPMC) [47] (**Figure 6A**). *Fam53C^-/-^* and *Fam53C^+/-^* mice were recovered at the expected frequency from *Fam53C^+/-^* crosses in the IMPC colony (with a low number of mice analyzed, **Figure S6A**), but with a significant trend towards fewer of the homozygous mutant mice in our colony at Stanford University (with more mice analyzed, **Figure 6B**). Based on the cell cycle arrest phenotypes observed in culture, *Fam53C^-/-^* may have been expected to have a reduced body size, similar for example to Cyclin D1 knockout mice [48, 49]. We observed a significantly lower body mass at weaning for males in our colony (**Figure 6C**). This phenotype may be dependent on the husbandry conditions as there were no significant differences in body weight between knockouts and wild-type controls, males or females, in the IMPC dataset (although again with only n=8 mice in each group there) (**Figure S6B**). Histological analysis of the brain and other tissues and organs in adult control and knockout mice did not reveal any gross defects (**Figure S6C**). We note that the IMPC behavioral analysis had shown significant differences between wild-type and *Fam53C^-/-^* male mice in a “latency to first transition into dark” test, in which the knockout mice showed a decreased exploration of a new environment, suggestive of possibly increased anxiety (n=6 *Fam53C^-/-^*males and n=7 *Fam53C^-/-^* females compared to historical controls) (**Figure 6D**). However, we did not repeat similar experiments in our colony. In a survival study, we did not detect any differences between controls and *Fam53C^-/-^*mice (**Figure S6D**). These observations in mice with minimal phenotypes linked to FAM53C loss are in stark contrast to the strong phenotypes observed in cells in culture and suggest mechanisms of compensation *in vivo* that remain to be identified.

**Figure 6:**
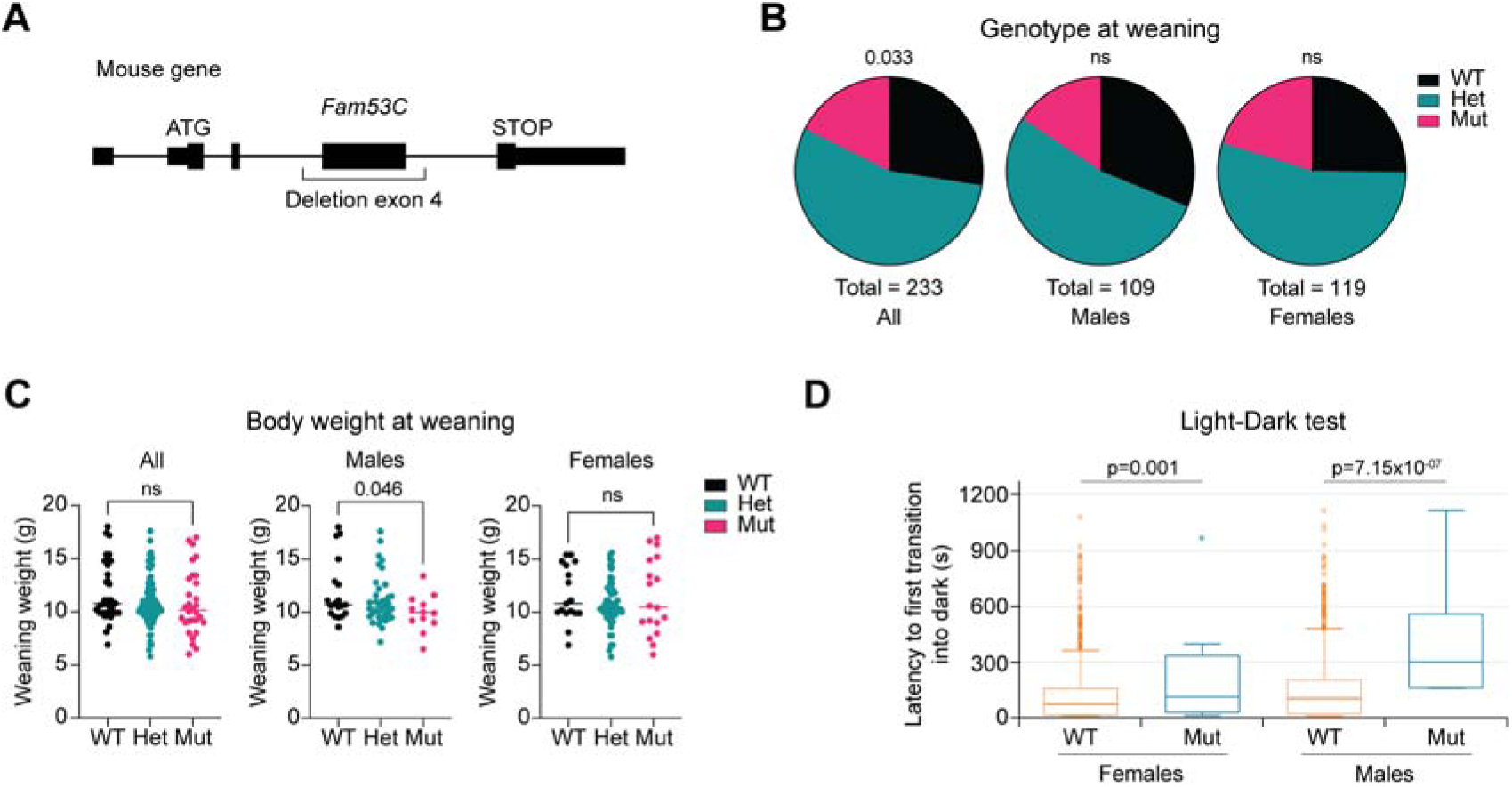
*Fam53C* knockout mice are viable and display limited phenotypes. **A.** Cartoon of the *Fam53C* mutant allele, with deletion of the major coding exon (exon 4) (not to scale). **B.** Genotypes of mouse pups at weaning from *Fam53C^+/-^* crosses at the Stanford facility. WT: wild-type; Het, heterozygous mutant mice; Mut, homozygous mutant mice. **C.** Body weight analysis at weaning for mice generated by *Fam53C^+/-^* crosses. **D.** Measure of the latency to first transition into a dark chamber (Light-Dark test) for control (n=1998 males and n=2037 females – including historical controls) and *Fam53C* knockout mice (n=6 males and n=7 females). Note the extremely different sizes in the cohorts of control and mutant mice, which may affect the results of the statistical analysis. P-values for (B) calculated by a Chi-squared test. P-values for (C) calculated by unpaired t tests. The details of the Linear Mixed Model framework for statistical analysis for (D) are available on the IMPC web site.

## DISCUSSION

Here we sought to identify novel regulators of the G1/S transition using the DepMap database as a starting point. Our analysis led us to focus on FAM53C, a factor which was previously uncharacterized as a cell cycle factor. Our data place FAM53C upstream of the CycD/CDK4,6-RB and p53-p21 pathways at the G1/S transition of the cell cycle. Our data further show that DYRK1A is a key partner of FAM53C in the control of cell cycle progression at this transition.

FAM53C is conserved throughout vertebrates, but little is known about the function of the FAM53 protein family. Previous work on FAM53B in medaka fish had shown a role in linking DNA-binding proteins to modulators of transcription in the control of proliferation [50]. Previous work on mammalian FAM53A had also suggested a connection with Bone Morphogenetic Protein/Transforming Growth Factor beta (BMP/TGFβ) signaling [51] and with p53 signaling [52], and our data connecting FAM53C loss to p53 activation may suggest an ancestral connection to p53 for this family.

DYRK1A controls the G1/S transition of the cell cycle [10, 12, 13, 41–43, 53, 54] but also plays a role in differentiation, especially in developing neurons [21, 55–59]. Thus, it is possible that FAM53C contributes to the control of cellular differentiation, including by regulating the cellular localization of DYRK1A [40], its kinase activity, or by other mechanisms. Because of the role of DYRK1A in the developing brain and in defects associated with Down syndrome [19–21], changes in FAM53C levels and/or activity may be expected to affect brain development. The behavioral data from the International Mouse Phenotyping Consortium (IPMC) support this idea, even though these experiments were performed with a small number of animals and will require future validation. In particular, the cellular basis of this behavioral phenotype, if repeated, will require a more in-depth analysis of the brain of mutant mice, possibly at different time points during development, as our histological analysis did not reveal any major changes in the brain of knockout mice compared to controls in adult animals. Notably, human *FAM53C* variants in public databases are linked to phenotypes such as height and Alzheimer’s disease (“co-localization analysis” from the Open Targets Genetics initiative [60]), as well as IGF1 serum levels (data from the Global Biobank Engine [61]). Disruption of IGF1 signaling is a candidate contributor to stunted growth and brain development defects in Down syndrome [62]. These observations provide further rationale to investigate FAM53C function *ex vivo* and *in vivo*, both in the regulation of cell cycle progression and in brain development and function.

Our data indicate that FAM53C controls DYRK1A kinase activity beyond its previously described ability to control DYRK1A cellular localization [40]. Our phosphorylation assays further suggest that DYRK1A may phosphorylate FAM53C, suggesting a role for FAM53C as a possible competitive inhibitor against other substrates. But It is also possible that this phosphorylation in turn affects FAM53C localization and/or activity. The identification and the mutation of the DYRK1A phosphorylation site(s) on FAM53C will be a first step to address this point in the future.

While our work identifies a FAM53C-DYRK1A-CycD/CDK4,6-RB pathway at the G1/S transition of the cell cycle, it is likely that FAM53C has other functions in cells. In particular, our AP-MS interactome data suggest that FAM53C may play additional roles later in the cell cycle, including during G2/M through associations with other kinases like PLK1 and AURKA. Whether these kinase interactions are inhibitory similar to the FAM53C-DYRK1A interaction remains to be determined. We did not observe cell cycle arrest at other stages of the cell cycle beyond G1, but future experiments, including in synchronized cell populations or by single cell tracking, will help address the role of FAM53C at other key cell cycle checkpoints.

With the approval of CDK4/6 inhibitors for the treatment of breast cancer and the development of a variety CDK inhibitors [63–65], it has become even more important than before to better understand the mechanisms regulating the activity of these kinases. Our data indicate that low levels of FAM53C may activate DYRK1A kinase activity towards Cyclin D, thereby decreasing CDK4/6 activity in cells. Our mass spectrometry analysis also identified CDK4 as a possible interactor of FAM53C. Although we did not pursue this observation, a possible direct interaction between FAM53C and CDK4 may also regulate CDK4 activity in cells. The identification of FAM53C as a regulator of the DYRK1A-CycD/CDK4,6-RB pathway suggests that strategies to decrease FAM53C levels in cells may help enhance the anti-tumor effects of CDK inhibitors.

## Methods

### Ethics statement

Mice were maintained at Stanford University’s Research Animal Facility according to practices prescribed by the NIH and by the Institutional Animal Care and Use Committee (IACUC) at Stanford University. Additional accreditation of Stanford University Research Animal Facility was provided by the Association for Assessment and Accreditation of Laboratory Animal Care (AAALAC). The study protocol was approved by the Administrative Panel on Laboratory Animal Care (APLAC) at Stanford University (protocol 13565).

### Animal studies

We imported *Fam53C* knockout mice from the International Mouse Phenotyping Consortium (IMPC [47], www.mousephenotype.org) (*Fam53^cem1(IMPC)J^* allele, C57BL/6NJ background). This allele was generated by electroporating the Cas9 protein along with 2 guide sequences 5’-ACTGCATTTTTGAGGAGGAG-3’ and 5’-GTTAAAATTCAATACTGCCA-3’, which resulted in a 1151 bp deletion beginning at Chromosome 18 position 34,767,928 bp and ending after 34,769,078 bp (GRCm38/mm10). This mutation deletes exon 4 and 366 bp of flanking intronic sequence including the splice acceptor and donor and is predicted to cause a change of amino acid sequence after residue 46 and early truncation 27 amino acids later. Genotyping was performed using the Mouse Direct PCR kit (for genotyping, B40015) following the protocol provided by the Jackson Laboratory (strain Stock No: 032892, C57BL/6NJ-*Fam53c^em1(IMPC)J/Mmjax^*) using a forward primer for the wild-type allele (5’-CCTGGGAACTCTTCTGTCTAGAGT-3’), a forward primer for the mutant allele (5’-TGGCATTACCACTTCACAGC -3’), and a common reverse primer (5’-CTAAGGACTAACTTGACAGGGCAGA-3’) (wild-type band: 99 bp; mutant band: 90 bp). We note that we have not been able to detect the FAM53C protein reliably with current antibodies in immunoblot on mouse cells, making it impossible to ascertain full knockout at the protein level.

Data generated by the IPMC were downloaded from their web site (https://www.mousephenotype.org/data/genes/MGI:1913556)[47]. For the cohort generated at Stanford to measure body weight and to assess phenotypes during aging, mice were generated from heterozygous crosses.

Histological analysis was performed on hematoxylin and eosin-stained paraffin sections prepared by HistoWiz.

### DepMap analysis

The DepMap analysis was performed directly from the DepMap portal in 2020 (20q2 datasets) (https://depmap.org/portal)[27].

### Cell culture and small molecules

Human cell lines were grown in DMEM high glucose medium supplemented with 10% bovine growth serum (BGS) (Fisher Scientific) and 1% penicillin-streptomycin-glutamine (Gibco). All cell lines tested negative for mycoplasma. RB and p53 mutant RPE-1 cells were a kind gift from Evgeny Zatulovskiy and Jan Skotheim [66]. Control and p53 knockout HCT116 cells were a kind gift from Mengxiong Wang and Laura Attardi.

The following small molecule inhibitors were used in cells: palbociclib HCl (Selleckchem #S1116) and puromycin (ThermoFisher #A1113803). For DYRK1A kinase inhibition, we used SM13797 from Biosplice [44] at a concentration of 1 µM with volume matched DMSO vehicle control unless otherwise noted.

Human induced pluripotent stem cells (hiPSCs) were cultured in Essential 8 medium (Thermo Fisher Scientific, A1517001) on cell culture plates coated with vitronectin (Thermo Fisher Scientific, 14190). They were passaged approximately every 6 days using 0.5 mM EDTA (Thermo Fisher Scientific, 15575) once they reached about 80% confluency. Cell integrity was confirmed using high-density SNP arrays, and routine PCR testing was performed to ensure cultures remained free of mycoplasma.

The differentiation of iPSCs into the three germ layers was performed using the STEMdiff™ Trilineage Differentiation Kit (STEMCELL Technologies, #05230).

### Knock-down, knock-out, and overexpression

FAM53C knock-down experiments in human cell lines were performed using 5 µM of ON-TARGETplus pooled siRNAs (Horizon – Dharmacon^TM^), and ON-TARGET Control #1 as negative controls. Cells were transfected using the Lipofectamine RNAiMAX reagent, following the manufacturer’s recommendations. Lentiviral vectors encoding HA-FAM53C (C-terminal tag) (VectorBuilder) or GFP control (a gift from the Artandi lab). Lentivirus was produced in 293T cells, and cells were infected with viral supernatant on two consecutive days, followed by four days of selection with puromycin.

hiPSCs (8 × 10^5^ cells) were electroporated in P3 primary cell nucleofector solution (Lonza, V4XP-3032) with 2 µg of FAM53C KO Plasmid containing GFP expression system (Santa Cruz Biotechnology, sc-412179) using a 4D-Nucleofector (program: DC100) (Lonza). After nucleofection, the cells were transferred to Matrigel-coated plates (Corning, 354277) and cultured in mTESR medium (STEMCELL Technologies, 100-0276) with 10 µM ROCK inhibitor (Y27632) at 37°C for 24 hr. GFP-positive cells were sorted using a FACSAria II flow cytometer (BD Biosciences) and cultured immediately in mTESR medium containing 10 µM Y27632 and Antibiotic-Antimycotic (Thermo Fisher Scientific,15240062). Approximately 300 GFP-positive cells were plated per well in Matrigel-coated six-well plates. Colonies were harvested in QuickExtract DNA Extraction Solution (LGC Biosearch Technologies, QE09050). Then, the genomic region around the CRISPR-Cas9 target site for *FAM53C* was amplified by PCR using Q5® Hot Start High-Fidelity 2X Master Mix (New England Biolabs, M0494S). To select the *FAM53C* knockout clone, we used the following primer pair: forward, 5′-GGCCCAAGATTCCTCTCGAC-3′, and reverse, 5′-GGGTCTCACCCCAGTTTCTC-3′. To examine the specific region of the knockout, RNA was extracted from the initially selected clones using the RNeasy Plus Mini Kit (Qiagen, 74136). cDNA synthesis was then performed using the SuperScript™ IV CellsDirect™ cDNA Synthesis Kit (Invitrogen, 11750150). The cDNA was amplified by PCR and subsequently sequenced. The primers used for this process were: forward, 5′-GCAGACTCTGGATGAGCTGAAATG-3′, and reverse, 5′-TCTCTTGGTGGCAGGGATACT-3′.c The sequencing results confirmed that the clone had a deletion of exon 3 and exon 4 (844 bp) (see gene map in Figure S5A). Potential off-target effects of the sgRNA were assessed using the RGEN tool (https://www.rgenome.net/), with PCR primers for detecting mutations at potential off-target sites (*ZFAND5* (forward, 5′-GCGGCGAGTGCGTTAGT-3′, and reverse, 5′-TTTGTTTCTCTGGGTCGTGGTG-3′), *MAP6D1* (forward, 5′-GGCTACTCGGACCTCGACA-3′, and reverse, 5′-GGTTCCAACTCGGCTGAAGG-3′), and *DCAF10* (forward, 5′-AAGTTTGGGTCAAGATCCTGGT-3′, and reverse, 5′-AAGGCCAAGTATACTCATAAGTGAGG - 3′) (see sequencing results in Figure S5B,C).

### Generation of hCOs from hiPSCs

The generation of human cortical organoids (hCOs) from hiPSCs followed a previously published protocol [67]. In brief, hiPSCs were dissociated into single cells using Accutase (Innovate Cell Technologies, AT-104) and seeded into AggreWell plates (STEMCELL Technologies, 34815) at a density of 3 × 10^6^ cells per well, using Essential 8 medium (Thermo Fisher Scientific, A1517001) supplemented with 10 µM Rock inhibitor Y-27632 (Selleckchem, S1049). The next day, spheres were collected and transferred to 10 cm ultra-low-attachment dishes (Corning, 3262). For the first 6 days, they were cultured in Essential 6 medium (Thermo Fisher Scientific, A1516401) with SB-431542 (10 µM, Tocris, 1614), dorsomorphin (2.5 µM Sigma-Aldrich, P5499), and XAV-939 (2.5 µM, Tocris, 3748), with daily medium changes. From Day 7, hCOs were maintained to Neurobasal A medium (Thermo Fisher Scientific, 10888) containing B27 (Thermo Fisher Scientific, 12587), along with 20 ng/ml epidermal growth factor (R&D Systems, 236-EG) and 20 ng/ml basic fibroblast growth factor (R&D Systems, 233-FB), with daily medium changes for 8 days, followed by changes every other day until Day 25. At day 25, the hCOs were switched to a Neurobasal A medium with B27, 20 ng/ml brain-derived neurotrophic factor (BDNF; Peprotech, 450-02), and 20 ng/ml NT3 (Peprotech, 450-03) for an additional 20 days, with medium changes every other day. From Day 45 onward, the hCOs were maintained in Neurobasal A medium with B27, with medium changes every 3-4 days.

### Quantitative immunoassay, immunoblot analysis, and immunofluorescence

For immunoassays with human cell lines, cells extracts were prepared in RIPA buffer with Roche cOmplete™ ULTRA proteasome and PhosSTOP™ phosphatase inhibitor cocktail tablets (Millipore Sigma #05892791001 and PHOSS-RO). Protein extracts were quantified using the Pierce™ BCA Protein Assay Kit according to manufacturer’s instructions (Thermo Fisher Scientific, 23227). For quantitative immunoassays, the capillary-based Simple Western^TM^ assay was performed on the Wes^TM^ system (ProteinSimple) according to the manufacturer’s protocol with 1 μg of protein used per lane. Compass software (ProteinSimple) was used for protein quantification, using the default settings unless otherwise specified. For immunoblot analysis, cell lysates were denatured in Laemmli buffer and boiled at 95°C for 5 minutes. 10 μg of protein was loaded into each lane unless otherwise specified. Samples were run on NuPAGE 4 to 12% Bis-Tris gels (Thermo Fisher Scientific # NP0322BOX) and transferred to nitrocellulose membranes using iBlot 2 transfer stacks (Thermo Fisher Scientific IB23002). Membranes were blocked in 10% milk in TBS-T (20 mM Tris, 150 mM NaCl, 0.1% Tween 20) for 1 hr. Primary antibodies were diluted in 5% milk in TBS-T and incubated overnight at 4°C. Secondary antibodies (Jackson ImmunoResearch anti-rabbit #111-035-144; anti-mouse #115-035-003) were diluted 1:10,000 in 5% milk in TBS-T and incubated for 1 hr at room temperature. Chemiluminescence was detected using Amersham ECL Prime Western Blotting Detection Reagent (GE Healthcare #RPN2236). The following primary antibodies were used for human cell lines: HSP90 (1:2000; Cell Signaling Technology, #4877), GAPDH (1:5000; Thermo Fisher, PA5-79289), FAM53C (1:250; Thermo Fisher Scientific, PA5-60125), HA tag (1:1000; Cell Signaling Technology, #3714), p21 (1:1000; Cell Signaling Technology, #2947), Cyclin D1 (1:500; Cell Signaling Technology, #2922), and DYRK1A (1:200; Cell Signaling Technology, #8765).

For experiments with hCOs, the organoids were lysed on ice using RIPA buffer with protease (Santa Cruz Biotechnology, sc-24948A) and phosphatase inhibitors (GenDEPOT, P3200-001) through gentle agitation, and incubated at 4°C for 1 hr. The lysates were then centrifuged at 14,000 × g for 15 minutes, and the supernatant was collected. Protein concentrations in the supernatant were measured using the Pierce™ BCA Protein Assay Kit (Thermo Fisher Scientific, 23225). The lysates were denatured with Bolt™ LDS Sample Buffer (Invitrogen, B0007) at 95°C for 5 minutes. Then, 7 µg of the samples were loaded onto Bolt™ 4-12% Bis-Tris Protein Gels (Invitrogen, NW04120BOX) and transferred to a PVDF membrane using the iBlot™ 2 Transfer Stacks and iBlot2 system (Method: P0) (Thermo Fisher Scientific, IB24002). The membranes were blocked with 5% BSA (GenDEPOT, A0100-005) in TBST (Tris-buffered saline with 0.1% Tween 20, Boston BioProducts, BM-301X) for 1 hr. They were then incubated overnight at 4°C with primary antibodies in TBST containing 5% BSA: DYRK1A (1:1000; Abcam, ab65220), FAM53C (1:1000; Thermo Fisher Scientific, PIPA5114093), CyclinD1 (1:1000; Cell Signaling, # 2922S), p-CyclinD1 (1:1000; Cell Signaling, #3300), P21 (1:1000; Cell signaling, # 2947S), and β-actin (1:1000; Cell signaling. #4970S). After washing with 5% TBST, the membranes were incubated with horseradish peroxidase-conjugated secondary antibodies at room temperature for 1 hr: Anti-mouse IgG (1:2000; Cell Signaling, #7076), and Anti-rabbit IgG (1:2000; Cell Signaling, #7074). The SuperSignal™ West Femto Maximum Sensitivity Substrate (Thermo Fisher Scientific, 34095) was used for signal development, and the iBright 1500 (Thermo Fisher Scientific, A44114) was used for protein band detection. Band intensities were quantified using ImageJ software (version 1.53t, NIMH, Bethesda, MD) with normalization to background and to the β-actin control.

For immunofluorescence analysis of differentiated iPSCs, cells were fixed in a 10% formalin solution (Sigma) at 4°C for 15 minutes, followed by washing with PBST (PBS with 0.1% Tween 20). Permeabilization was carried out using PBS containing 0.1% Triton X-100 (Sigma) for 20 minutes. Then, the samples were incubated in 3% BSA for 1 hr before being treated overnight at 4°C with the respective primary antibodies : PAX6 for ectoderm (1:300; BioLegend, PRB-278P), Brachyury for mesoderm (1:300; R&D Systems, AF2085-SP), and SOX17 for endoderm (1:300; Cell Signaling, 81778S). The cells were then rinsed with PBST and incubated with Alexa Fluor-conjugated secondary antibodies (1:500; 488 or 594; Thermo Fisher Scientific) for 1 hr at room temperature. Samples were observed using a Zeiss LSM 980 confocal microscope (Carl Zeiss).

### Cell cycle and cell death analyses

Cells were counted using a Countess 3 instrument on default settings (Thermo Fisher Scientific), and dead cells were excluded using 0.4% Trypan Blue solution (Thermo Fisher Scientific, T10282). For palbociclib treatment (with DMSO as a control), cells were treated with 0.5 µM for 24 hr. Cell cycle analysis was performed using BrdU pulsed for 2 hr (10 µg/mL, Calbiochem #203806) and propidium iodide (PI) staining[7] unless otherwise indicated. For cell death assays, Annexin V/PI staining was performed and analyzed as described before[7]. Flow cytometry was performed on a BD FACSAria II (BD Biosciences) and data was collected using BD FACSDiva. Data were analyzed using Cytobank Community software (Beckman Coulter Life Sciences).

For experiments with hCOs, the EdU assays were performed using the Click-iT™ EdU Alexa Fluor™ 647 Flow Cytometry Assay Kit (Thermo Fisher Scientific, C10424), following the manufacturer’s instructions. Briefly, hCOs were incubated with 10 µM EdU for 24 hr, then dissociated using Accutase and resuspended in staining buffer with 3% BSA and 0.5 mM EDTA. After fixation and permeabilization, the cells were incubated with the Click-iT™ reaction cocktail for 30 minutes at room temperature, protected from light. DAPI was added for DNA staining, and the cells were analyzed using a FACSAria II flow cytometer (BD Biosciences) and FCS Express 7 software (DeNovo Software).

### Bulk RNA sequencing and analysis

For FAM53C knock-down experiments in RPE-1 cells, 1×10^6^ control and siRNA-treated samples were collected in duplicate 48 hours post-transfection, and RNA was isolated using the RNeasy Plus Micro kit (Qiagen #74034). Library preparation and sequencing were performed by Novogene using the Illumina NextSeq 500 platform to obtain ∼20 million paired reads/sample. Raw sequencing reads were trimmed using CutAdapt (v2.10)[68] with the TruSeq sequencing adapter (5’-AGATCGGAAGAGCACACGTCTGAACTCCAGTCAC-3’), and a minimum read length of at least 25. Reads were then aligned to the UCSC hg38 genome with HiSat2 [69] using reverse strandedness and discarding unaligned reads. Counts were assigned to genes (hg38 GTF) using featureCounts [70] Downstream differential expression analysis was performed using the Deseq2 package [71]. GO term enrichment of differentially expressed genes was performed using ClusterProfiler [72].

### FAM53C interactome analysis

#### Tandem affinity purification

10 mL packed cell volume of HEK-293T cells expressing LAP-tagged proteins [73] were re-suspended with 40 mL of LAP-resuspension buffer (300 mM KCl, 50 mM HEPES-KOH [pH 7.4], 1 mM EGTA, 1 mM MgCl2, 10% glycerol, 0.5 mM DTT, protease inhibitor [A32965, Thermo Fisher Scientific]), lysed by gradually adding 1200 µL 10% NP-40 to a final concentration of 0.3%, then incubated on ice for 10 min. The lysate was first centrifuged at 14,000 rpm (27,000 g) at 4°C for 10 min, and the resulting supernatant was centrifuged at 43,000 rpm (100,000 g) for 1 hr at 4°C to further clarify the lysate. High speed supernatant was mixed with 500 µL of GFP-coupled beads [73] and rotated for 1 hr at 4°C to capture GFP-tagged proteins, then washed five times with 1 mL LAP200N buffer (200 mM KCl, 50 mM HEPES-KOH [pH 7.4], 1 mM EGTA, 1 mM MgCl2, 10% glycerol, protease inhibitors, and 0.05% NP40). After re-suspending the beads with 1 mL LAP200N buffer lacking protease inhibitors, the GFP tag was cleaved by adding 40 µg PreScission-protease and rotating tubes at 4°C for 16 hr. PreScission-protease eluted supernatant was added to 100 µL of S-protein agarose (69704-3, EMD Millipore) to capture S-tagged protein. After washing three times with LAP200N buffer and twice with LAP0 buffer (50 mM HEPES-KOH [pH 7.4], 1 mM EGTA, 1 mM MgCl2, and 10% glycerol), purified protein complexes were eluted with 50 µL of 2X LDS buffer and boiled at 95°C for 3 min. Samples were then run on Bolt Bis-Tris Plus Gels (NW04120BOX, Thermo Fisher Scientific) in Bolt MES SDS Running Buffer (B0002, Thermo Fisher Scientific). Gels were fixed and stained according to Colloidal Blue Staining Kit (LC6025, Thermo Fisher Scientific) with Optima LC/MS grade water (W6-4, Fisher Scientific) at room temperature. The buffer was then replaced with Optima water prior to cutting the bands into eight pieces. The gel slices were then destained, reduced, and alkylated followed by in-gel digestion using (200 ng) Trypsin/LysC (V5073, Promega) as previously [74]. Tryptic peptides were extracted from the gel bands and dried in a speed vac. Prior to LC-MS, each sample was reconstituted in 0.1% formic acid, 2% acetonitrile, and water.

#### Chemicals and Reagents

The reagents used in this analysis were obtained from various sources. From Sigma-Aldrich, we obtained 2-Chloroacetamide (CAM; cat. no. C0267), and Potassium hydroxide (KOH; cat. no. P5958). Trifluoroacetic acid (TFA; cat. no. AAL06374AC), Acetic acid (AcOH; cat. no. A11350), Optima™ LC/MS Grade Water (cat. no. W6-4), and 99.5% Formic acid, LC/MS Grade (FA; cat. no. A117-50) were supplied by Fisher Scientific. Tris(2-carboxyethyl)phosphine hydrochloride (TCEP-HCl; cat. no. PG82080) and Halt™ Protease and Phosphatase Inhibitor Cocktails, EDTA-Free (cat. nos. 78425 and 78428, respectively), were purchased from Thermo Fisher Scientific. Honeywell provided the LC/MS grade acetonitrile (ACN; cat. no. 14261-1L). Additionally, Trypsin/Lys-C Mix, Mass Spec Grade (cat. no. V5073) from Promega, and Empore C18 47 mm Extraction Disk (cat. no. 320907D) from Empore were used. Liquid chromatography was conducted using a Bruker PepSep C18 10 cm packed column with 1.5 µm beads and 150 µm I.D (cat. No. 1893483) attached to a ZDV Sprayer with 20 µm I.D (cat. No. 1865710).

#### Stage-Tips Clean Up

Homemade Stage-Tips were constructed using two C18 Empore disks, following established procedures. The fabricated Stage-Tips were subjected to a washing process, which involved two washes with 100 µL of methanol, one wash with 100 µL of 80% acetonitrile/0.1% acetic acid, and two washes with 100 µL of 1% acetic acid. Acidified peptides were loaded onto the Stage-Tips. Subsequently, the Stage-Tips were washed three times with 100 µL of 1% acetic acid to remove salts. Finally, the peptides were eluted from the Stage-Tips using two elution steps of 30 µL each, with 60% acetonitrile/0.1% acetic acid as the elution buffer.

#### Liquid Chromatography Setup

A nanoELute ultra-high-pressure nano-flow chromatography system was utilized and directly coupled online with a hybrid trapped ion mobility spectrometry—quadrupole time-of-flight mass spectrometer (timsTOF Pro, Bruker) employing a nano-electrospray ion source (CaptiveSpray, Bruker Daltonics).

#### Chromatographic Conditions

The liquid chromatography was conducted at a constant temperature of 50°C, employing a reversed-phase column (PepSep column, 10 cm × 150 µm i.d., packed with 1.5 µm C18-coated porous silica beads, Bruker) connected to the 10 µm emitter (Bruker). The mobile phase consisted of two components: Mobile Phase A, comprising water with 0.1/2% formic acid/ACN (v/v), and Mobile Phase B, comprising ACN with 0.1% formic acid (v/v).

#### Gradient Elution

Peptide separation was achieved using a linear gradient from 2-33% Mobile Phase B within 60 min. This was followed by a washing step with 95% Mobile Phase B and subsequent re-equilibration. The chromatographic process maintained the flow rate at 400 nL/min.

#### MS Acquisition

Samples were analyzed using the timsTOF HT Mass Spectrometer in DDA-PASEF mode. The TIMS elution voltage was calibrated linearly to obtain reduced ion mobility coefficients (1/K0) by using three selected ions from the Agilent ESI-L Tuning Mix (m/z 622, 922, 1222). The mass and ion mobility ranges were set from 100 to 1700 m/z and 0.7 to 1.3 1/K0, respectively. Both ramp and acquisition times were set at 100 ms. Precursor ions suitable for PASEF-MS/MS were chosen from TIMS-MS survey scans using the PASEF scheduling algorithm. A polygon filter was applied to the m/z and ion mobility plane to prioritize features likely representing peptide precursors over singly charged background ions. The quadrupole isolation width was set to 2 Th for m/z < 700 and 3 Th for m/z > 700, with collision energy linearly increased from 20 to 60 eV as ion mobility ranged from 0.6 to 1.6 (1/K0).

#### Silver staining

5 µL of samples containing LDS buffer and DTT prepared for tandem affinity purification and mass spectrometry described above were mixed with 1.25 µL of 500 mM iodoacetamide (0210035105, MP Biomedicals). Proteins were separated in a 4–12% Bis-Tris gel (NP0321BOX, Invitrogen), followed by fixation of the gel overnight in 50% methanol at room temperature. The gel was impregnate with solution C (0.8% (w/v) silver nitrate (S6506, SIGMA), 207.2mM ammonium hydroxide (A6899, SIGMA) and 18.9 mM sodium hydroxide) for 15 min, followed by rinsing with water twice. The image was then developed in solution D (0.005% citric acid and 0.0185% formaldehyde in Milli-Q) until intensity of the bands increase to optimal level. The reaction was then terminated by adding stop solution (45% methanol and 10% acetic acid).

#### Data Analysis

Raw mass spec result files were processed using Bionic (Protein Metrics, Inc.) software (version: 4.5.2) with the following parameters: precursor mass tolerance: 20 ppm, fragment mass tolerance: 40 ppm; fragmentation type: QTOF/HCD; carbamidomethyl as Cys fixed modification, variable modifications of Met oxidation, Asn and Gln deamidation, pyro-Glu formation at N-term Glu and Gln, N-term Acetylation and Ser, Thr and Tyr phosphorylation, with maximum 3 variable modification; trypsin with maximum 2 missed cleavages and fully specific mode; identifications were filtered at 0.01 FDR at the protein level; a fasta library of all human refseq proteins (curated and predicted) was used that was downloaded on 2021/07/20.

Spectral counts were normalized to NSAF values [75] and significance of enrichment of bait-association was calculated as described previously [76].

### Proteomic analysis

BioGRID [77]. Gene ontology (GO) term enrichment analysis and protein-protein interaction analysis was performed using Metascape with the default settings in the “Express Analysis” function. We used Enrichr [78] to analyze the top candidate interactors from the IP/MS experiments (66 proteins, p-val<0.05) on 07/25/2024.

### Recombinant protein expression and purification

Recombinant proteins were expressed in *Escherichia coli* BL21 (DE3) cells. The cells were transformed with plasmids encoding the target protein fused with either a His-tag (FAM53C) or a GST-tag (DYRK1A). Transformed BL21 cells were cultured in LB medium supplemented with 100μg/mL Ampicillin. Protein expression was induced at an OD₆₀₀ of 0.8 using 1 mM IPTG. The cultures were grown overnight at 20°C.

The bacterial cells were harvested by centrifugation and resuspended in lysis buffer containing 50 mM Tris-HCl pH 8.0, 150 mM NaCl, and 1 mM TCEP. The cells were lysed by cell homogenizer and clarified by centrifugation. For His-tagged proteins, the lysate was applied to a nickel-nitrilotriacetic acid (Ni-NTA) column pre-equilibrated with equilibration buffer (50 mM Tris-HCl pH 8.0, 150 mM NaCl, 1mM TCEP, 10 mM imidazole). After incubation for 30 min, resin with bound protein was washed with the same buffer but containing 30 mM imidazole and subsequently bound proteins were eluted with 200 mM imidazole buffer. For GST-tagged proteins, the lysate was applied to a GST-affinity resin pre-equilibrated with equilibration buffer (50 mM Tris-HCl pH 8.0, 150 mM NaCl, 1 mM TCEP), and the protein was eluted using the same buffer but containing 20 mM reduced glutathione.

The His or GST tag was cleaved by incubating the eluted protein with 0.1 mg/mL GST-TEV protease overnight at 4°C. Following cleavage, the protein mixture was passed through a GST-affinity column to remove GST-TEV and cleaved tag. The tag-free protein was further purified by ion exchange chromatography using a Q Sepharose column. The protein solution was loaded onto the Q column pre-equilibrated with 50 mM Tris-HCl pH 8.0 and 50 mM NaCl. Bound proteins were eluted using a gradient of 50 mM to 1 M NaCl. As a final purification step, the protein was subjected to size exclusion chromatography on a Superdex 75 column equilibrated with buffer containing 50 mM Tris-HCl pH 8.0, 500 mM NaCl, and 1 mM TCEP. Fractions containing the target protein were pooled and concentrated using centrifugal filtration. The purity of the final protein preparation was confirmed by SDS-PAGE and protein stored at −80°C in 10% glycerol.

To obtain biotinylated DYRK1A, we used sortase labeling as described [79], utilizing the N-terminal glycine that is left following TEV cleavage. 50 µM DYRK1A was reacted with 500 µM of a synthetic biotin-LPETGG peptide and a final concentration of 20 µM purified His-tagged sortase in a final reaction volume of 500 µL. The reaction was incubated overnight at 4°C. Following the reaction, the DYRK1A was purified again by passing over Ni^2+^-NTA and Superdex 75 columns and was stored as above. The Cyclin D1 used in the recombinant protein kinase assay is a complex of CDK4-CycD1 purified as previously described [80].

### Kinase assays

Protein samples were mixed in a buffer containing 25 mM Tris-HCl pH 8.0, 150 mM NaCl, 20 mM MgCl₂, and 1 mM DTT. A mixture of non-radioactive and [γ-P^32^] ATP (∼10 µCi) was added to the assay samples at a final concentration of 200 nM and allowed to react for 15 or 30 min as indicated. The samples were subjected to SDS-PAGE for protein separation. The SDS-PAGE was performed on a 4-20% gradient at 200V for 40 min under denaturing conditions. Following electrophoresis, the gel was dried down completely to capture the signal from the radiolabeled ATP. The radioactive decay was detected by exposing the dried gel to a phosphor screen overnight. The phosphor screen was scanned using a GE Typhoon Trio Imager.

### SM13797 Selectivity Profile

Biochemical IC_50_ values for CLK1-CLK4, DYRK1A, CDK1 and GSK3β were determined by acoustically transferring SM13797 to 384-well plates (Echo 550, LabCyte) and by performing kinase assays using ThermoFisher LanthaScreen platform for CLK4 or Z’LYTE™ platform for the other kinases following the manufacturer’s instructions. IC_50_ values were calculated from 11-point dose response curves using Dotmatics Studies software. The full kinome screen (484 kinases) was performed using ThermoFisher SelectScreen profiling service with SM13797 at 1μM. The IC_50_ values were subsequently determined for each kinase demonstrating > 90% inhibition. Target engagement IC_50_ values were determined using the Promega NanoBRET® TE Intracellular Kinase Assay platform in transiently transfected HEK293T cells. IC_50_ values were determined from 10-point dose response curves using Dotmatics Studies software.

### Biolayer Interferometry

BLI experiments were performed using an eight-channel Octet-RED96e (Santorius). Experiments were performed in an assay buffer containing 25 mM Tris pH 8.0, 150 mM NaCl, 1 mM DTT, 2 mg/mL BSA, and 0.2% (v/v) Tween. Samples were formatted in a 96-well plate with each well containing 200 µL. For each experiment, we used eight streptavidin biosensor tips (Santorius) that were loaded with the 200 nM biotinylated DYRK1A and dipped into FAM53C analyte at varied concentrations. All experiments were accompanied by reference measurements using unloaded streptavidin tips dipped into the same analyte-containing wells. Experiments in which analyte concentration was varied also contained a zero-analyte reference. Data were processed and fit using Octet software version 7 (Santorius). Before fitting, all datasets were double reference-subtracted, aligned on the *y* axis through their respective baselines, aligned for interstep correction through their respective dissociation steps, and smoothened using Savitzy –Golay filtering. The first 20 s of association and 20 s of dissociation were used with a 1:1 binding model. The association and dissociation rate constants as well as standard errors were averaged across the set of sensorgrams. The equilibrium dissociation constant K_D_ was obtained from a steady state analysis of all concentrations of analyte fit to the equation Response = (R_max_ * [Analyte])/(K_D_ + [Analyte]).

### Statistical analysis

Statistical significance was assessed using Prism GraphPad software, unless otherwise stated in the methods above. The tests used are noted in the figure legends and accompanying Supplementary Tables. Data are represented as mean ± standard deviation unless otherwise stated. Statistical significance was assessed using Prism GraphPad software, unless otherwise stated in the methods above. The tests used are noted in the figure legends and accompanying Supplementary Tables. Data are represented as mean ± standard deviation unless otherwise stated.

### Data availability

Sequencing data from RNA sequencing are available from the Gene Expression Omnibus (GEO) under accession numbers GSE282945. The mass spectrometry proteomics data have been deposited to the ProteomeXchange Consortium via the PRIDE [81] repository with the dataset identifier PXD055829 and 10.6019/PXD055829. Data related to the IPMC genotyping and phenotyping of *Fam53C* mutant mice is available on the IPMC web site. All other data are available in the article and supplementary materials, or from the corresponding author upon reasonable request.

## Acknowledgements

We thank all the members of the Sage lab for their help and support throughout this study. Research reported in this publication was supported by the NIH (J.S., S.M.R., J.M.S., P.K.J. P01CA254867). J.S. is the Elaine and John Chambers Professor in Pediatric Cancer.

---

**Figure S1 related to Figure 1:**
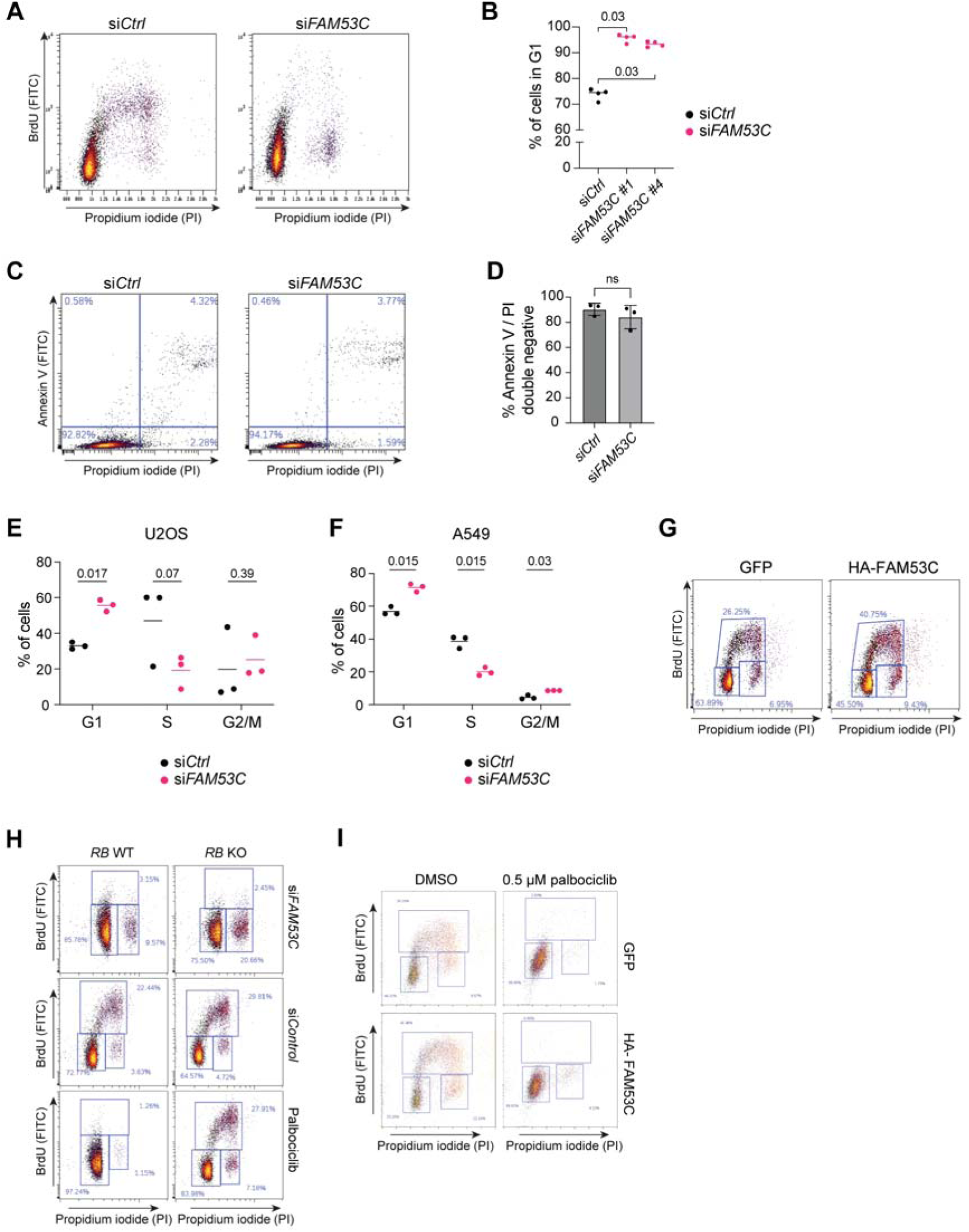
Identification of FAM53C as a positive regulator of cell cycle progression in G1. **A.** Representative example of flow cytometry analysis for BrdU/PI staining of RPE-1 cells with control siRNAs and FAM53C siRNAs (related to Figure 1E). **B.** Fraction of cells in G1 from BrdU/PI staining in RPE-1 cells, with or without FAM53C knock-down, using two individual siRNAs (N=3). **C.** Representative example of flow cytometry analysis for Annexin V/PI staining of RPE-1 cells with control siRNAs and FAM53C siRNAs. The double negative cells are the live cells. **D.** Quantification of (C) (N=3). **E.** Cell cycle analysis by BrdU/PI staining in control and FAM53C knock-down U2OS cells (N=3). **F.** Cell cycle analysis by BrdU/PI staining in control and FAM53C knock-down A549 cells (N=3). **G.** Representative example of flow cytometry analysis for BrdU/PI staining of RPE-1 cells with GFP or FAM53C overexpression. **H.** Representative example of flow cytometry analysis for BrdU/PI staining of RPE-1 cells in control and RB knockout RPE-1 cells as in Figure 1J. **I.** Representative example of flow cytometry analysis for BrdU/PI staining of RPE-1 cells in control and FAM53C-overexpressing RPE-1 cells as in Figure 1K. P-values for (B), (D), (E), and (F) were calculated by t-test.

**Figure S2, related to Figure 2:**
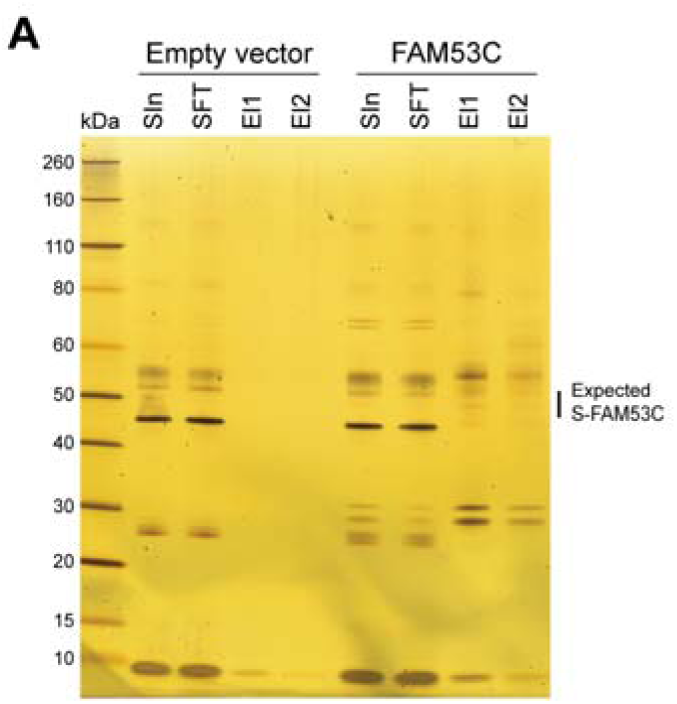
The FAM53C interactome identifies cell cycle factors. |**A.** Silver stain of protein fractions in the AP/MS experiment. Sin: S-bead input (cleavage of protein from the GFP-beads before binding to s-beads); SFT: S-bead flowthrough (solution after binding to S-beads). El1, El2: elution 1 and elution 2 from S-beads.

**Figure S3, related to Figure 3:**
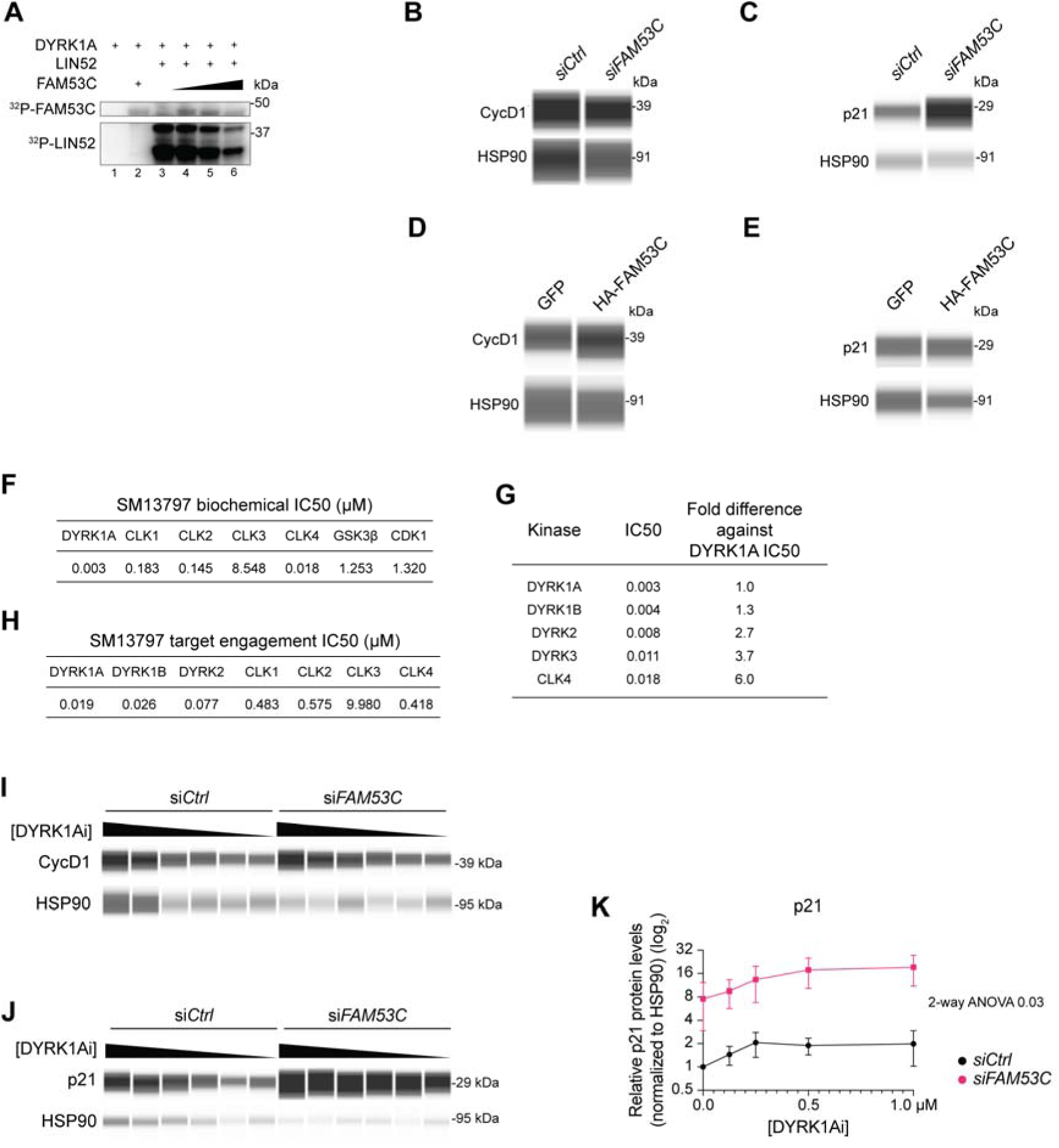
FAM53C can inhibit DYRK1A function in cells. **A.** Recombinantly expressed LIN52 (10 µM) was phosphorylated by DYRK1A (400 nM) for 30 min alone or in the presence of increasing amounts of FAM53C. FAM53C concentration was 1.6, 5, 15 µM (lanes #4-6). Lane #1 contains DYRK1A without substrate. Representative experiment of N=2 experiments. **B.** Representative immunoassay for Cyclin D1 protein levels relative to the loading control HSP90 in FAM53C knock-down (si*FAM53C*) RPE-1 cells compared to controls (si*Ctrl*) at 24 h (Related to Figure 3B). **C.** Representative immunoassay for p21 protein levels relative to the loading control HSP90 in FAM53C knock-down (si*FAM53C*) RPE-1 cells compared to controls (si*Ctrl*) at 24 h (Related to Figure 3C). **D.** Representative immunoassay for Cyclin D1 protein levels relative to the loading control HSP90 in RPE-1 cells expressing HA-FAM53C compared to GFP controls (Related to Figure 3D). **E.** Representative immunoassay for p21 protein levels relative to the loading control HSP90 in RPE-1 cells expressing HA-FAM53C compared to GFP controls (Related to Figure 3E). **F.** SM13797 biochemical IC_50_ values determined using the ThermoFisher LanthaScreen platform for CLK4 or Z’LYTE™ platform for the other kinases. **G.** List of kinases inhibited at 90% or more by SM13797 (1 µM) using ThermoFisher SelectScreen service and showing IC_50_ within 20-fold of that of DYRK1A. IC_50_ determination for DYRK1B, DYRK2 and DYRK3 were performed by ThermoFisher SelectScreen service. **H.** Cellular target engagement profile of SM13797 against CLK/DYRK family members. Target engagement assay IC_50_ values were determined using the Promega NanoBRET® TE Intracellular Kinase Assay platform in transiently transfected HEK293T cells. **I.** Representative immunoassay for Cyclin D1 protein levels in RPE-1 cells treated with or without different concentration of the SM13797 DYRK1Ai (related to Figure 3F). **J.** Representative immunoassay for p21 protein levels in RPE-1 cells treated with or without different concentration of the SM13797 DYRK1Ai. **K.** Quantification of immunoassays for p21 protein levels in RPE-1 cells treated with or without different concentration of the SM13797 DYRK1Ai, as in (I) (N=3 per concentration). P-value for (K) was calculated by 2-way ANOVA.

**Figure S4, related to Figure 4:**
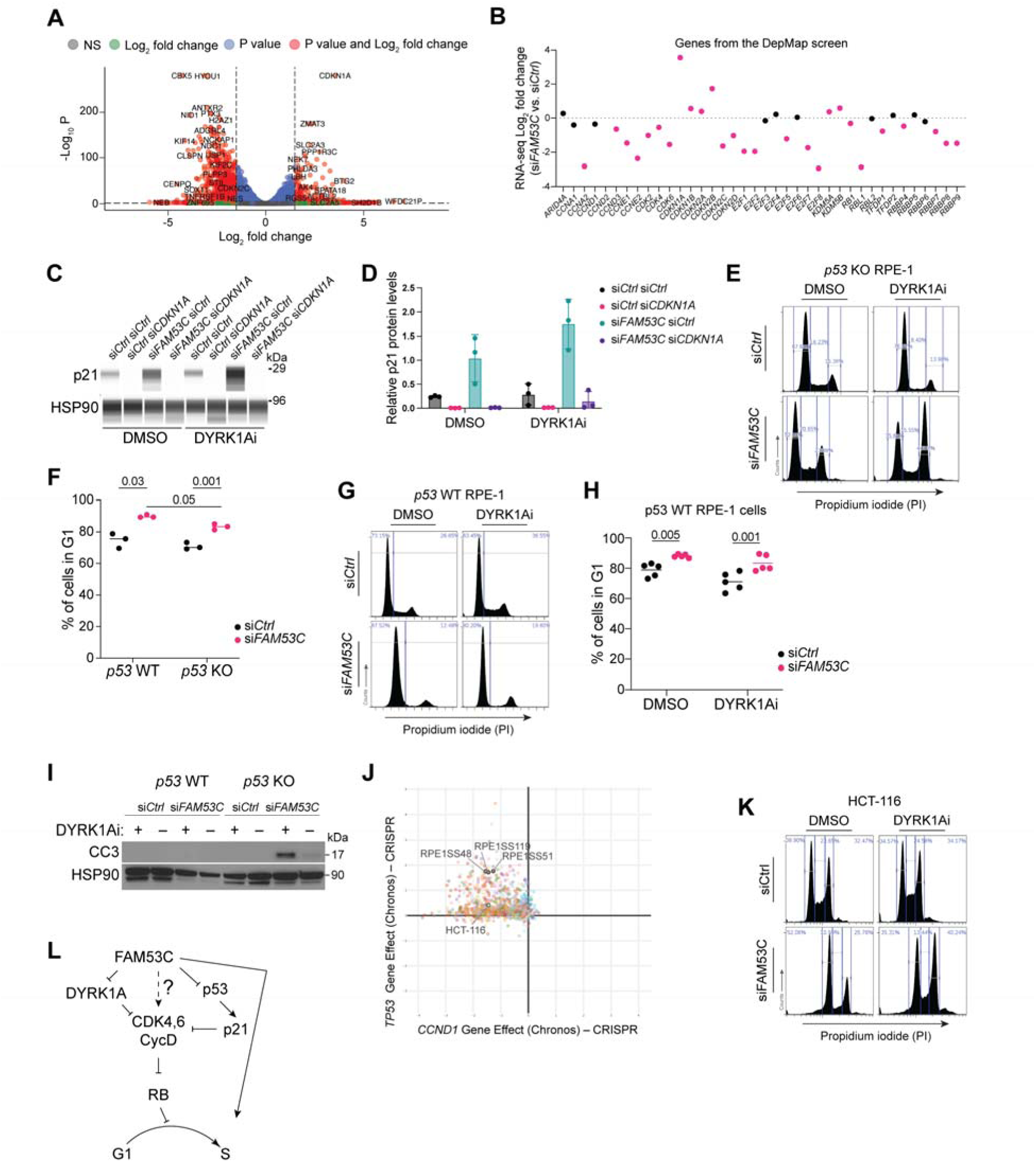
FAM53C knock-down activates p53. **A.** Volcano plot from the RNA-seq data comparing control and FAM53C knock-down RPE-1 cells (48 h after siRNA transfection). See Supplementary Table S5. **B.** Fold-change analysis of the 38 genes selected in the initial DepMap screen (Supplementary Table S1) in FAM53C knock-down cells compared to controls (from the RNA-seq data). Pink dots: P-value<0.05. **C.** Immunoassay for p21 (encoded by *CDKN2A*) in RPE-1 cells treated with DMSO or the DYRK1Ai, with or without FAM53C knock-down (48 hours of knock-down and treatment). HSP90 serves as a loading control (N=3). **D.** Quantification of (C) (N=3). **E.** Representative example of flow cytometry analysis for PI staining of *p53* knockout (KO) RPE-1 cells treated with DMSO or the DYRK1Ai, with or without FAM53C knock-down. **F.** Fraction of cells in G1 in control (wild-type, WT) and *p53* knockout (KO) RPE-1 cells, with or without FAM53C knock-down (N=3), as in (E). Note that a G1 arrest is still present in *p53* knockout cells with the FAM53C knock-down. **G.** Representative example of flow cytometry analysis for PI staining of *p53* wild-type (WT) RPE-1 cells treated with DMSO or the DYRK1Ai, with or without FAM53C knock-down. **H.** Fraction of cells in G1 in *p53* WT RPE-1 cells as in (G) (N=5). **I.** Immunoblot for cleaved caspase 3 (CC3) in *p53* wild-type and knockout RPE-1 cells, with or without FAM53C knock-down, and with or without DYRK1Ai treatment. HSP90 serves as a loading control (N=1). **I.** Fraction of cells in G1 in RPE-1 as in (H). Note that cells arrest in G1 upon FAM53C knock-down and that this arrest is not rescued by the p21 knock-down (ns, not significant). **J.** Correlation for the gene effects of *CCND1* and *Tp53* (coding for Cyclin D1 and p53, respectively) in the DepMap CRISPR dataset. Four cell lines are highlighted, three derivatives of RPE-1 cells and HCT-116. **K.** Representative example of flow cytometry analysis for PI staining of HCT-116 cells, with or without FAM53C knock-down, with or without DYRK1Ai treatment, as in Figure 4F. **L.** Cartoon placing FAM53C in the G1/S transition of the cell cycle upstream of the RB and p53 pathways. P-values for (F) and (H) were calculated by paired t-test.

**Figure S5, related to Figure 5:**
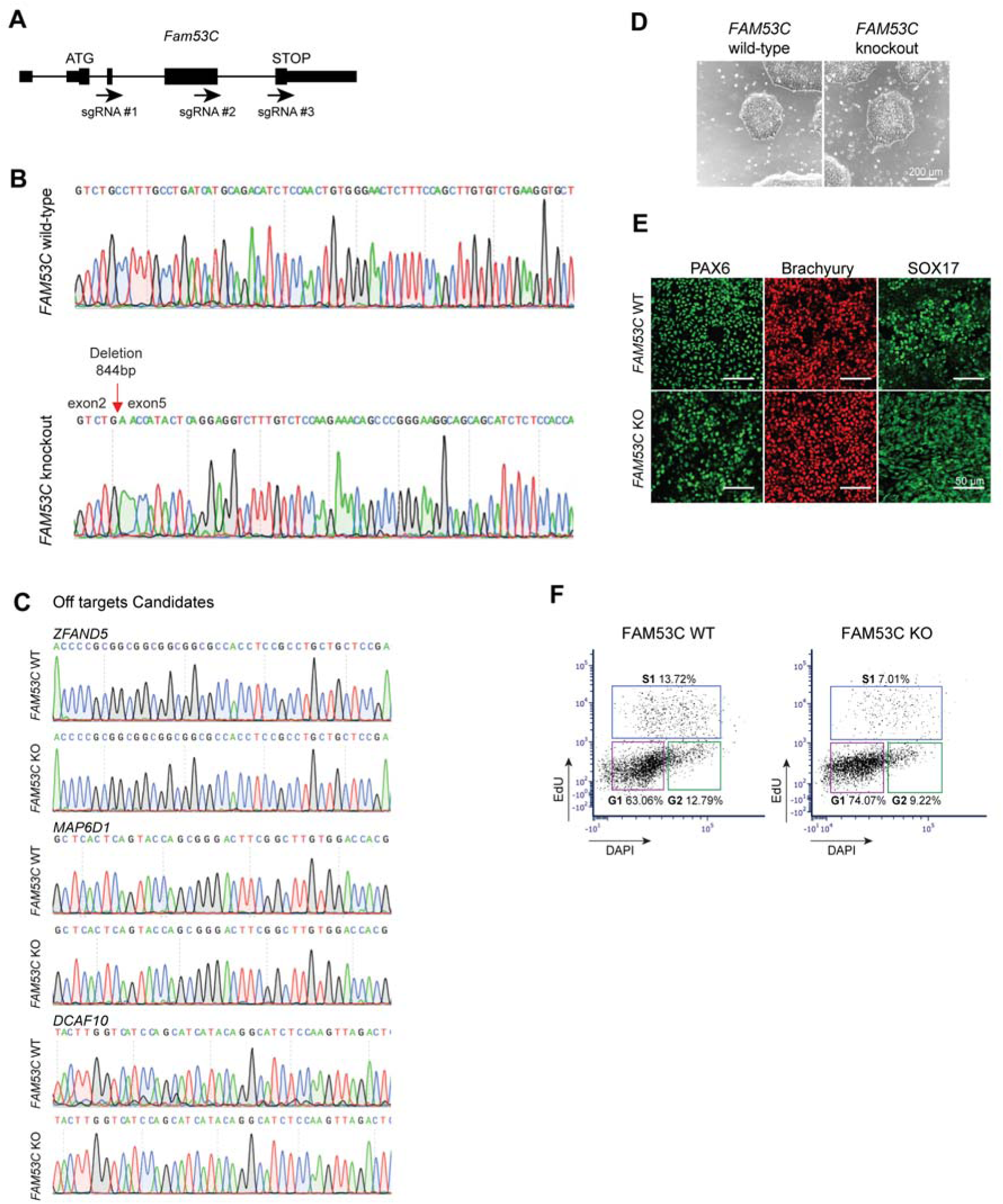
Loss of FAM53C impairs the development of human cortical organoids. **A.** Cartoon of the *FAM53C* human gene, with the location of the sgRNAs used to knockout the gene in iPSCs (not to scale). **B.** Sequence histogram showing gene inactivation upon CRISPR/Cas9-mediated deletion. **C.** Sequence histogram showing no changes in the sequence of possible off target genes. **D.** Representative images of wild-type and knockout iPSC colonies. Scale bar, 200 µm. **E.** Representative images of immunofluorescence staining of pluripotency markers in wild-type and knockout iPSCs. Scale bar, 50 µm. **F.** Representative example of flow cytometry analysis for EdU/DAPI staining of hCOs, as in Figure 4E.

**Figure S6, related to Figure 6:**
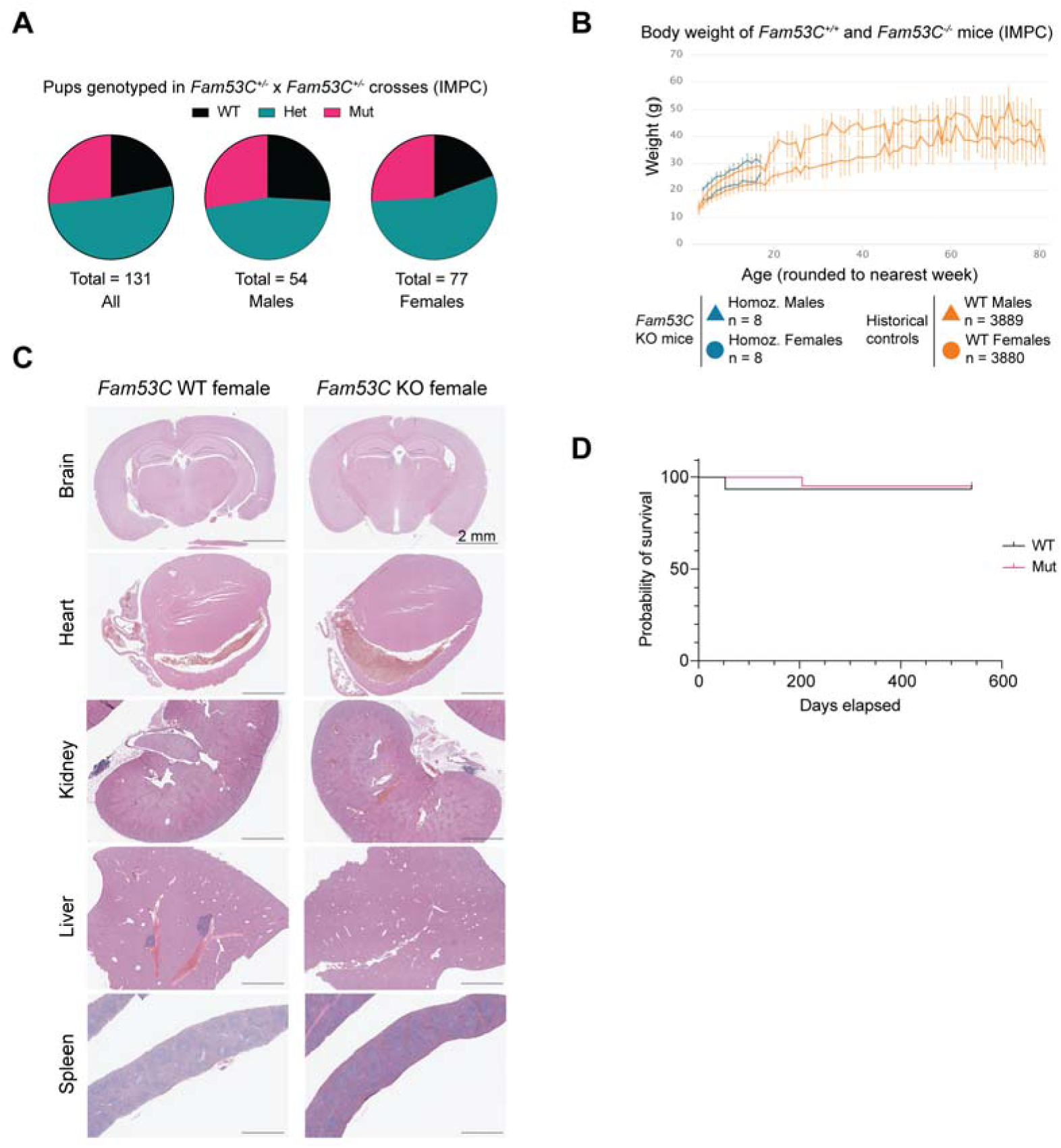
*Fam53C* knockout mice are viable and display limited phenotypes. **A.** Genotypes of mouse pups at weaning from *Fam53C^+/-^* crosses at the IMPC facility. **B.** Body weight analysis of a large cohort of historical controls and *Fam53C^-/-^* mice. **C.** Histological analyses (sections stained with hematoxylin and eosin, H&E) for the indicated tissues from mice generated by *Fam53C^+/-^* crosses. Representative images for one wild-type and one knockout female are shown out of two females and two males analyzed (age, 18 months). Scale bar, 2 mm. **D.** Survival analysis for N=23 *Fam53C^-/-^* mice (7 males and 15 females) and N=16 wild-type controls (7 males and 9 females) from heterozygous crosses. The analysis was stopped at 18 months.

